# Single-domain antibodies represent novel alternatives to monoclonal antibodies as targeting agents against the human papillomavirus 16 E6 protein

**DOI:** 10.1101/388884

**Authors:** Melissa Togtema, Greg Hussack, Guillem Dayer, Megan Teghtmeyer, Shalini Raphael, Jamshid Tanha, Ingeborg Zehbe

**Author notes:** Corresponding Author for single-domain antibody-related inquiries. Corresponding Author for HPV-related inquiries. Email Addresses: Melissa Togtema Dr. Greg Hussack Dr. Guillem Dayer Megan Teghtmeyer Shalini Raphael Dr. Jamshid Tanha Dr. Ingeborg Zehbe.

## Abstract

Approximately one-fifth of all malignancies worldwide are etiologically-associated with a persistent viral or bacterial infection. Thus, there is particular interest in therapeutic molecules which utilize components of a natural immune response to specifically inhibit oncogenic microbial proteins, as it is anticipated they will elicit fewer off-target effects than conventional treatments. This concept has been explored in the context of human papillomavirus type 16 (HPV16)-related cancers, through the development of monoclonal antibodies and fragments thereof against the viral E6 oncoprotein. However, challenges related to the biology of E6 as well as the functional properties of the antibodies themselves appear to have precluded their clinical translation. In this study, we attempted to address these issues by exploring the utility of the variable domains of camelid heavy-chain-only antibodies (denoted as VHHs). Through the construction and panning of two llama immune VHH phage display libraries, a pool of potential VHHs was isolated. The interactions of these VHHs with recombinant E6 protein were further characterized using ELISA, Western blotting under both denaturing and native conditions, as well as surface plasmon resonance, and three antibodies were identified that bound recombinant E6 with affinities in the nanomolar range. Our results now lead the way for subsequent studies into the ability of these novel molecules to inhibit HPV16-infected cells *in vitro* and *in vivo*.

## 1. Introduction

Approximately 20% of cancers worldwide are etiologically-associated with persistent infection by microbes such as viruses (Feng *et al*. 2008, zur Hausen *et al*. 2009). Advantageously, when prophylactic measures are unavailable or ineffective, the opportunity remains to specifically target infected cells using therapeutic molecules which block the expression or function of oncogenic microbial proteins. As these proteins often bear minimal homology to those of the host, such molecules are anticipated to have fewer off-target effects on surrounding uninfected cells compared to conventional treatments, reducing treatment-associated toxicity and facilitating disease intervention at the earliest stages of lesion development (Hoppe-Seyler *et al*. 2018).

One of the most ubiquitous tumour viruses in humans is the papillomavirus (HPV). Due to its common transmission through sexual contact, almost all individuals will become infected with this keratinocyte-tropic DNA virus throughout their lifetime (zur Hausen 1996, WHO 2016). Twelve high-risk mucosal HPV types have been well-established as the causative agents in almost all cases of cervical cancer (Bouvard *et al*. 2009) and have been implicated in anogenital as well as oropharyngeal cancers (Crow 2012). Of these, HPV16 is the most commonly identified (Crow 2012, Mehanna *et al*. 2013). Most strikingly, the incidence of HPV-related oropharyngeal cancers in males has been on the rise in Canada, the United States, and Europe, outpacing that of HPV-related cervical cancers (Giuliano *et al*. 2015, Canadian Cancer Society 2016, Centers for Disease Control and Prevention, Division of Cancer Prevention and Control 2017). Of the products encoded by HPV16’s episomal genome (Figure 1A) (Doorbar *et al*. 2015), the E6 oncoprotein (~18 kDa) (Figure 1B) promotes the immortalization and malignant transformation of persistently infected cells through a variety of interactions with host intracellular proteins (Klingelhutz and Roman 2012, Vande Pol and Klingelhutz 2013), including degradation of the tumour suppressor protein p53 (Martinez-Zapien *et al*. 2016). As E6 is expressed in both precancerous lesions and tumours, and its inhibition can promote apoptosis when expression of the pro-proliferative E7 oncoprotein is retained, it has accordingly been recognized as an optimal HPV16-specific therapeutic target (Hoppe-Seyler *et al*. 2018).

**Figure 1.**
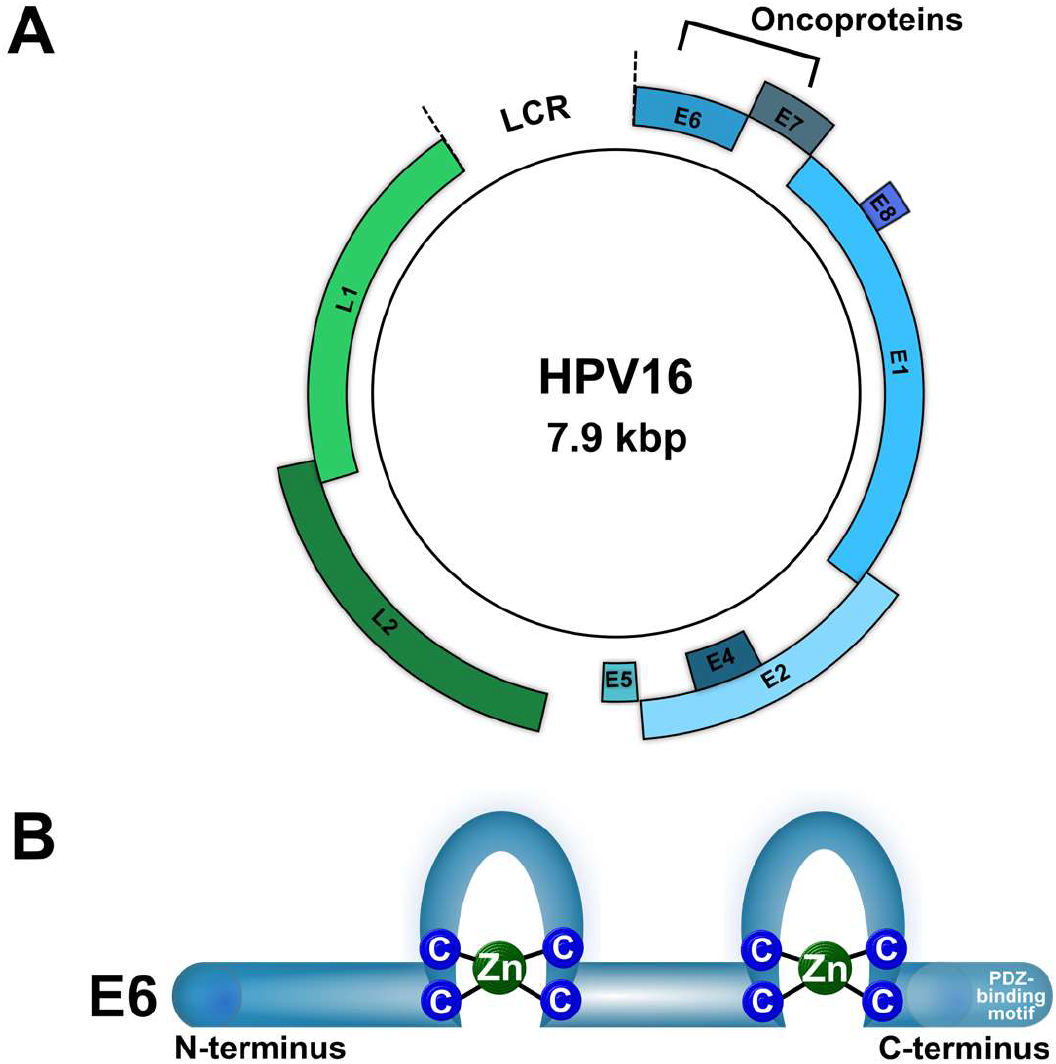
The HPV16 genome (~7.9 kbp) and E6 oncoprotein (~18 kDa). ***(A)** The circular, double-stranded DNA genome consists of a long control region (LCR) which regulates viral transcription, as well as seven early (E1, E2, E4, E5, E6, E7, and E8) and two late (L1 and L2) genes. The proteins encoded by the early genes promote viral persistence, replication, and release, whereas those encoded by the late genes create the viral capsid (Doorbar et al. 2015, Dreer et al. 2017, Harden and Munger 2017). The HPV16 genome (GenBank Accession #: K02718.1) was visualized using the Pathogen-Host Analysis Tool (Gibb et al. 2017). **(B)** The E6 oncoprotein contains two zinc-binding domains. It interacts with host proteins after first complexing with the hijacked ubiquitin ligase E6AP or via its C-terminal PDZ-binding motif, driving cancerous changes in infected cells (Klingelhutz and Roman 2012, Vande Pol and Klingelhutz 2013)*.

Inhibitory molecules of a transient nature may present fewer ethical concerns than those intended to permanently edit the viral genome (*e.g*., CRISPR/Cas9 (Stone *et al*. 2016)). They can be designed to target either E6 mRNA, preventing its translation, or the E6 protein itself, sterically hindering its intracellular interactions. Molecules which mimic components of a natural immune response are of particular interest as they utilize endogenous cellular pathways and are hypothesized to have high functionality with low toxicity (Togtema *et al*. 2012, Togtema *et al*. 2018). However, transcript silencing using synthetic small interfering RNAs (siRNAs) has been limited by challenges in achieving the desired therapeutic effects at clinically relevant concentrations, off-target effects, as well as siRNA stability and uptake *in vivo* (Togtema *et al*. 2018). Alternatively, monoclonal antibodies (mAbs) benefit from greater therapeutic specificity and longer intracellular half-lives than siRNA (Cao and Heng 2005). Several mAbs specific to the N-terminal region (clones 1F1, 6F4, 4C6) or the second zinc-binding domain (clones 1F5, 3B8, 3F8) of the HPV16 E6 protein have been isolated from immunized mice (Giovane *et al*. 1999, Masson *et al*. 2003, Lagrange *et al*. 2005) (Figure 2A). Preliminary studies by both us and others have demonstrated that, when transiently transfected into HPV16-positive cell cultures, these mAbs elicited a notable restoration of p53 protein levels (Courtête *et al*. 2007, Togtema *et al*. 2012) and that conjugation of the mAbs to a nuclear localization signal (NLS) improved their ability to access E6’s mainly nuclear location, further enhancing this response (Freund *et al*. 2013, Postupalenko *et al*. 2014). Nevertheless, the anticipated induction of apoptosis was not observed. HPV16 E6-specific single-chain variable fragments (scFvs) (i.e., antibody variable heavy (VH) and light (VL) chain domains joined together by a synthetic linker peptide) which are amenable to ectopic expression inside mammalian cells as intrabodies as well as to passive nuclear diffusion have also been explored (Sibler *et al*. 2003, Griffin *et al*. 2006, Lagrange *et al*. 2007, Amici *et al*. 2016) (Figure 2A). But because non-specific effects in HPV-negative cells as well as unexpected downstream cellular responses were observed (Lagrange *et al*. 2007, Amici *et al*. 2016), further research into their application is needed.

**Figure 2.**
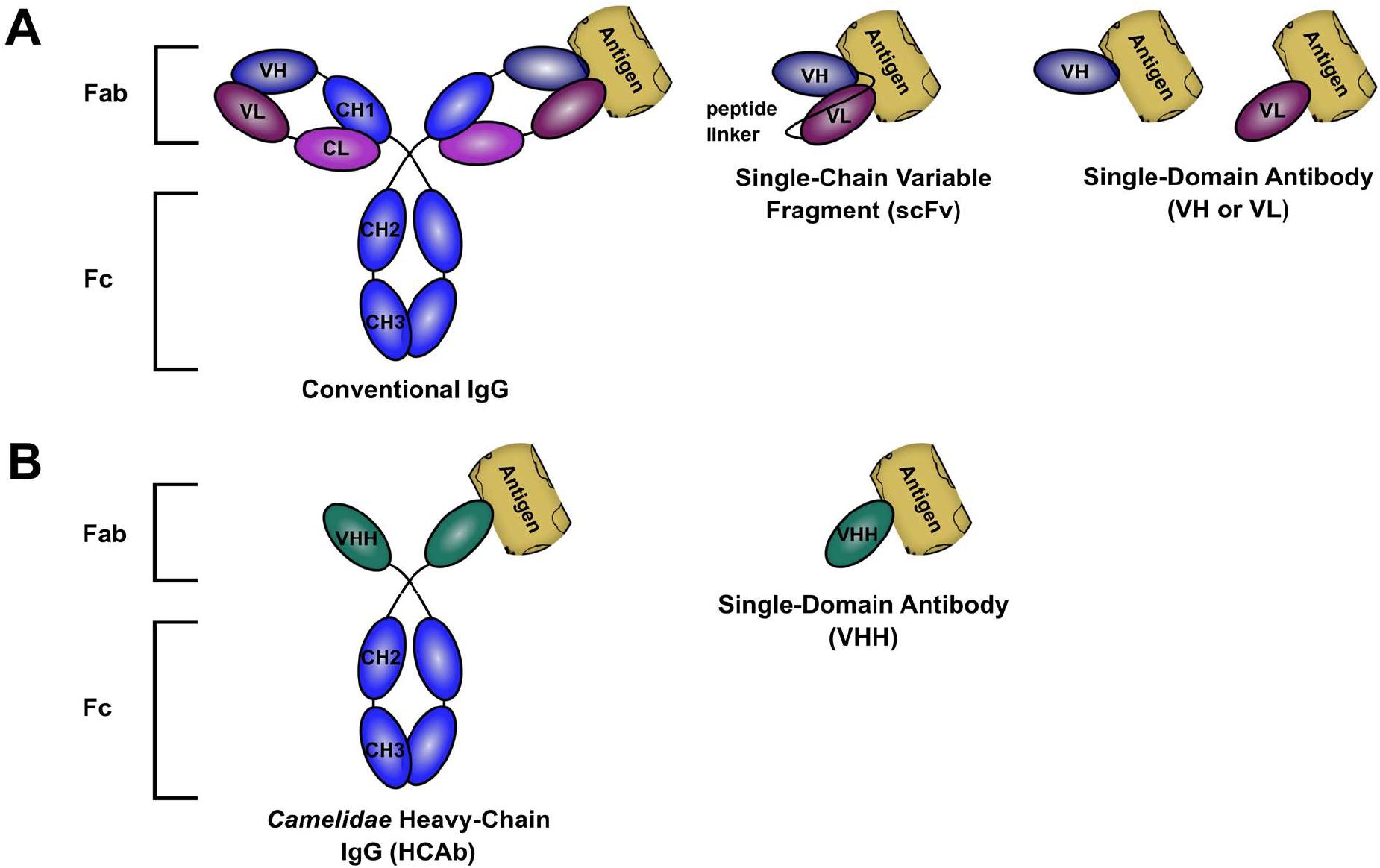
Conventional versus Camelidae heavy-chain IgGs. ***(A)** Conventional IgGs consist of two identical heavy and two identical light protein chains. The fragments of the antibody responsible for antigen binding (Fab) are each comprised of the variable domain and first constant domain of one heavy chain (VH and CH1) plus the variable domain and constant domain of one light chain (VL and CL). The paratope created by the VH and the VL together gives the antibody its specificity for a particular antigen. The second and third constant domains of the heavy chains (CH2 and CH3) form the fragment crystallizable (Fc) region by which the antibody interacts with other components of the immune system. Accordingly, smaller recombinant molecules such as single-chain variable fragments (scFvs: VH and VL plus linker) can retain the complete ability of the full-size antibody to interact with its antigen. These structural details were summarized from Janeway et al. (2001) and Muyldermans (2013). **(B)** Camelidae heavy-chain IgGs (HCAbs) consist of only two identical heavy chains. The CH1 domain is absent and interaction with the specific target antigen occurs solely through the variable domain of the heavy chain (VHH). Thus, VHH single-domain antibodies can retain the complete ability of the full-size antibody to interact with its antigen. These structural details were summarized from Muyldermans (2001), Flajnik et al. (2011), and Muyldermans (2013)*.

While mAb-based therapeutic molecules have been clinically-approved over the last few decades for various malignancies (Shepard *et al*. 2017), similar translation for HPV16-related cancers has not occurred, owing to challenging biological aspects of the HPV16 E6 protein itself, including its primarily nuclear location and potentially limited target epitope availability, as well as to technical hurdles in producing optimally-functional antibodies or fragments thereof. Instead, antibody fragments derived from unconventional sources, such as IgGs produced by *Camelidae* species (*e.g.*, llamas) which were discovered to naturally lack both light chains and CH1 domains (heavy-chain-only antibodies: HCAbs) in a subset of their antibody repertoire (Hamers-Casterman *et al*. 1993) (Figure 2B), may present useful, unexplored options. The variable domains of these HCAbs (denoted hereon as VHHs) can be isolated as single-domain antibodies, which conveniently retain the complete ability of the full-size antibody to interact with its antigen and demonstrate affinities for target antigens similar to those of conventional antibodies (Harmsen and De Haard 2007, Steeland *et al*. 2016) (Figure 2B). Due to their small size (~15 kDa) and the hydrophilic amino acid substitutions which evolved at the absent VL interface (Flajnik *et al*. 2011, Helma *et al*. 2015), VHHs possess several properties which may prove beneficial for E6-targeting including convex paratopes which can interact with antigen epitopes inaccessible to conventional mAbs and scFvs (Lauwereys *et al*. 1998, Stijlemans *et al*. 2004, De Genst *et al*. 2006, Flajnik *et al*. 2011, Steeland *et al*. 2016), robust thermal and chemical stability (Dumoulin *et al*. 2002, Muyldermans 2013), as well as the ability to facilely enter the nucleus through nuclear pores (Sibler *et al*. 2003). With superior solubility compared to conventional mAb fragments, VHHs are particularly amenable to both high-yield periplasmic expression in *E. coli* followed by transfection of the purified molecules into HPV-infected cells as well as to direct intrabody expression (Revets et al. 2005, Steeland et al. 2016, Böldicke et al. 2017). Lastly, VHHs also share a greater sequence homology to human than murine VHs, minimizing the extent of humanization required for clinical translation (Vu *et al*. 1997, Vincke *et al*. 2009).

New therapeutic applications for VHHs are continuously being identified (Steeland *et al*. 2016, Van Audenhove and Gettemans 2016), and several VHHs against targets involved in inflammatory/auto-immune, bone, neurological, hematological, oncological, and infectious diseases have already progressed to clinical trial evaluation (Steeland *et al*. 2016). Accordingly, upon recognition of their utility in the context outlined above, we sought to add to this growing compendium by isolating VHHs against the HPV16 E6 protein. Following the construction and panning of two llama immune VHH phage display libraries, phage ELISA indicated a pool of potential binders. The interactions of these VHH clones with recombinant E6 protein were then characterized using soluble ELISA, Western blotting under both denaturing and native conditions, as well as surface plasmon resonance (SPR) binding assays. Based on our findings, we hypothesize that subsequent studies will elucidate the ability of these novel molecules to inhibit HPV16-infected cells in *vitro* and in vivo.

## 2. Materials and Methods

### 2.1 Recombinant proteins

Three types of recombinant HPV16 E6 protein were used in this study (Figure 3). The first (denoted His6-GenScript E6) corresponded to the protein variant found in the cervical carcinoma-derived HPV16-positive cell line CaSki, which contains R10G and L83V amino acid substitutions from the reference sequence (GenBank Accession #: K02718.1) (Pattillo *et al*. 1977, Meissner *et al*. 1999, Zehbe *et al*. 2009). It was expressed with an N-terminal polyhistidine (His6) tag in E. *coli*, solubilized from inclusion bodies, purified using immobilized metal-affinity chromatography (IMAC), and dialyzed into buffer containing 1x PBS, pH 7.4, 1% sodium lauroyl sarcosine by GenScript (Piscataway, NJ, USA).

**Figure 3.**
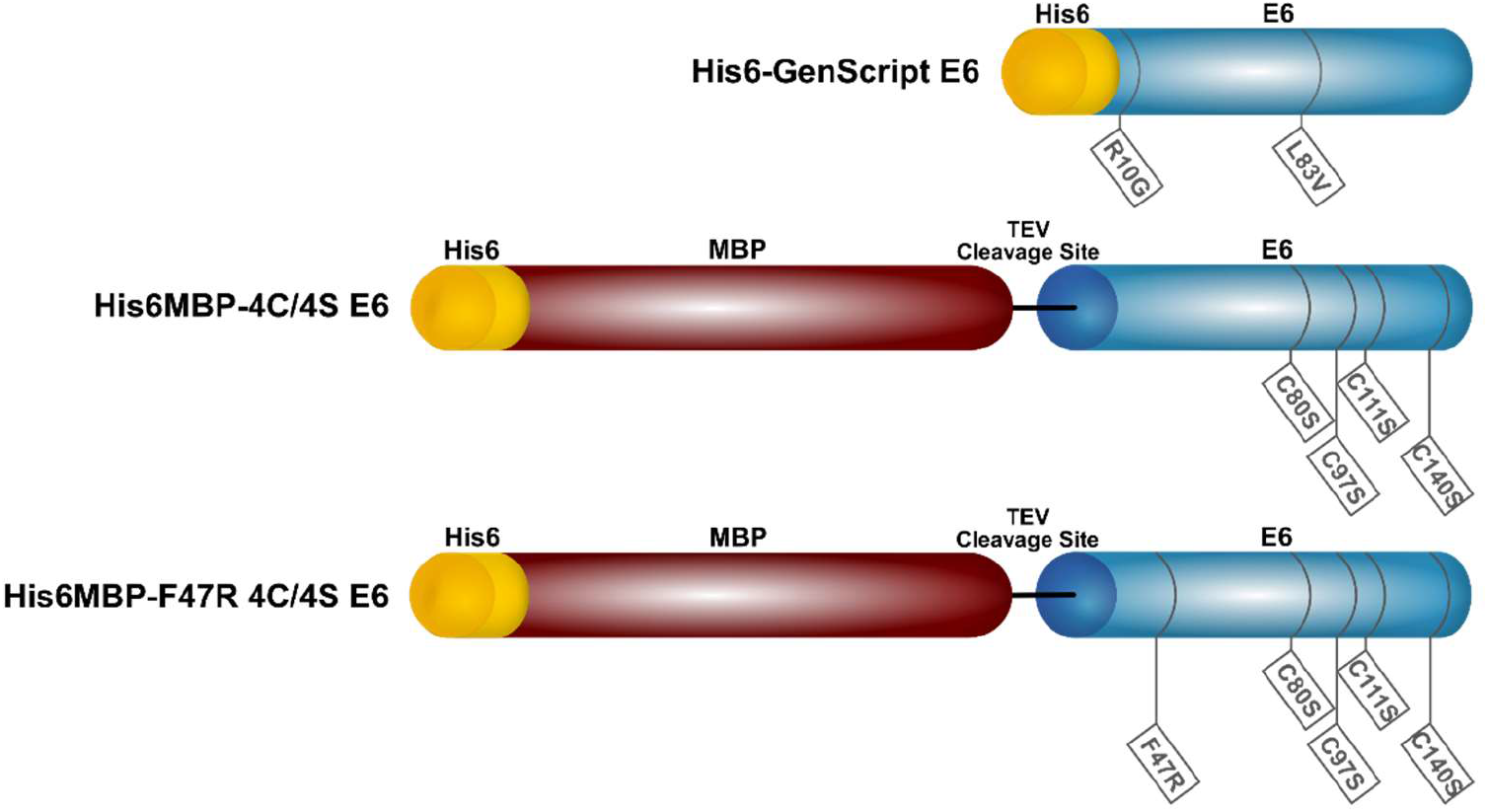
Schematic representation of the recombinant HPV16 E6 proteins used in this study. Their amino acid differences from the reference sequence (GenBank Accession #: K02718.1) as well as their respective polyhistidine (His6) tags and maltose binding protein (MBP) fusion partners are indicated.

The other two recombinant HPV16 E6 proteins (denoted His6MBP-4C/4S E6 and His6MBP-F47R 4C/4S E6) contained the solubility-enhancing amino acid substitutions C80S, C97S, C111S, C140S or F47R, C80S, C97S, C111S, C140S, respectively, and were expressed by us from the previously described pETM-41 plasmids (kindly provided by Dr. Gilles Travé, IGBMC, France) with an N-terminal His6-maltose binding protein (MBP) tag followed by a TEV protease cleavage site (Zanier *et al*. 2012, Zanier *et al*. 2013, Ramirez *et al*. 2015, Martinez-Zapien *et al*. 2016). Briefly, expression of each His6MBP-E6 protein was induced with 1 mM IPTG overnight at 15 °C in ClearColi® BL21 (DE3) *E. coli* (Lucigen; Middleton, WI, USA; Cat. #: 60810-1) grown in LB media. The bacterial pellets were resuspended in ice-cold lysis buffer (50 mM Tris-HCl pH 8.0, 25 mM NaCl) and stored for 1 h at −80 °C or overnight at −20 °C. SigmaFAST™ EDTA-Free Protease Inhibitor Cocktail Tablets (Sigma-Aldrich; Oakville, ON, Canada; Cat. #: S8830) and 2 mM DTT were then added, and the thawed cells lysed using 150 μg/mL lysozyme. The lysate was further incubated with 60 units/mL DNase I and then clarified by centrifugation. Finally, the recombinant antigen was purified from the clarified lysate using IMAC and dialyzed into buffer containing 50 mM Tris-HCl pH 6.8, 400 mM NaCl, 2 mM DTT. Recombinant MBP in 20 mM Tris pH 8.0 was purchased from Abnova (Walnut, CA, USA; Cat. #: P4989). All proteins were kept at −80 °C for long-term storage.

### 2.2 Llama immunizations

Two llamas were immunized through services available at Cedarlane Laboratories (Burlington, ON, Canada), using the “short schedule” similarly described by Baral *et al*. (2013). The first (Immunization #1) was injected with 100 μg of His6-GenScript E6 + Freund’s complete adjuvant on Day 0, followed by booster injections of 100 μg His6-GenScript E6 + Freund’s incomplete adjuvant on Days 21, 28 and 35. The second (Immunization #2) was injected with 250 μg of His6MBP-4C/4S E6 + 250 μg of His6MBP-F47R 4C/4S E6 + Freund’s complete adjuvant on Day 0, followed by booster injections of 250 μg His6MBP-4C/4S E6 + 250 μg His6MBP-F47R 4C/4S E6 + Freund’s incomplete adjuvant on Days 21, 28 and 35. From both llamas, pre-immune blood was collected prior to the initial immunization as well as blood samples on Days 35 and 42, to allow confirmation of a successful immune response and to provide the lymphocytes subsequently used in the preparation of the VHH phage display libraries, as described below.

### 2.3 Serology

Confirmation of a successful immune response was first obtained by screening the total serum from blood drawn on Days 0 (pre-immune), 35 and 42 for antibodies which reacted with the recombinant E6 proteins by enzyme-linked immunosorbent assay (ELISA), as similarly described by Hussack *et al*. (2011), Hussack *et al*. (2012a), and Baral *et al*. (2013). Briefly, 1 μg of His6-GenScript E6 diluted in PBS, or 0.5 μg His6MBP-4C/4S or His6MBP-F47R 4C/4S E6 diluted in storage buffer (50 mM Tris-HCl pH 6.8, 400 mM NaCl, 2 mM DTT) (100 μL/well) was coated in Nunc™ MaxiSorp™ 96-well plates (VWR; Mississauga, ON, Canada; Cat. #: CA10761-500) overnight at 4 °C. The wells were blocked with 5% (w/v) milk powder in PBS-T (PBS+0.05% (v/v) Tween 20) at 37 °C, prior to the application of serial serum dilutions for 1 h at room temperature. The wells were then washed with PBS-T and incubated with a goat anti-llama IgG + horseradish peroxidase (HRP) secondary antibody (Cedarlane Laboratories; Cat. #: A160-100P) diluted 1:10 000 in PBS for 1 h at room temperature. The wells were again washed with PBS-T and then incubated with 100 μL/well TMB substrate (Mandel Scientific; Guelph, ON, Canada; Cat. #: KP-50-76-00) for approximately 5 min. The reaction was stopped by the addition of 100 μL/well 1 M phosphoric acid and the absorbance read at 450 nm.

Day 0 and Day 42 total serum was then fractionated using protein A and protein G affinity chromatography as described by Hussack *et al*. (2012a) and Baral *et al*. (2013), with the addition of a second, pH 2.7 glycine buffer elution from the protein A column. Reducing SDS-PAGE was used to analyze the eluted G1 (IgG3 HCAb), G2 (IgG1 conventional antibody), A1 (IgG2a HCAb) and A2 (IgG2b/c HCAb) serum fractions, prior to confirmation of a positive HCAb immune response using ELISA, as similarly described above. In this instance, wells were coated with either 0.75 μg of His6-GenScript E6 diluted in PBS or a mix of 0.5 μg His6MBP-4C/4S E6 + 0.5 μg His6MBP-F47R 4C/4S E6 diluted in PBS (100 μL/well).

### 2.4 Construction of VHH phage display libraries

Two VHH phage display libraries were constructed, one corresponding to each immunization (i.e., Library #1: His6-GenScript E6 immunization and Library #2: His6MBP-E6 immunization). For each library, RNA was extracted from ~1.0 × 10^8^ frozen Day 42 lymphocytes using the PureLink™ RNA Mini Kit (Thermo Fisher Scientific; Mississauga, ON, Canada; Cat. #: 12183018A) and first-strand cDNA synthesis was performed using Superscript™ VILO™ Master Mix (Thermo Fisher Scientific; Cat. #: 11755050) with ~2 μg template RNA per reaction (total number of reactions: 4). PCR amplification of the VHH DNA was then completed, using the primers and procedures similarly described by Baral *et al*. (2013). Briefly, cDNA was first amplified in 50 μL reactions consisting of 3 μL cDNA, 5 pmol MJ1-3 primer mixture, 5 pmol of either CH2FORTA4 or CH2B3-F primer, 1 μL 10 mM dNTPs, 0.5 μL Platinum™ *Taq* DNA Polymerase (Thermo Fisher Scientific; Cat. #: 10966026), 1.5 μL *Taq* kit 50 mM MgCl2, 5 μL *Taq* kit 10x buffer, and 38 μL nuclease free H2O using thermocycler parameters of 94 °C for 5 min, 35 cycles of 95 °C for 30 s, 56 °C for 45 s and 72 °C for 60 s, and a final extension of 72 °C for 10 min. A total of 8 PCR reactions were performed and pooled for each primer pair. PCR products of ~600 bp (for reactions containing CH2FORTA4 primer) and ~650 bp (for reactions containing CH2B3-F primer) were extracted from 1.5% agarose gels, purified and then re-amplified in 50 μL reactions consisting of ~20 ng template DNA, 10 pmol MJ7 primer, 10 pmol of MJ8 primer, 1 μL 10 mM dNTPs, 0.5 μL Platinum™ *Taq* DNA Polymerase, 1.5 μL *Taq* kit 50 mM MgCl2, 5 μL *Taq* kit 10x buffer, and nuclease free H2O using thermocycler parameters of 94 °C for 5 min, 35 cycles of 95 °C for 30 s, 58 °C for 45 s and 72 °C for 60 s, and a final extension of 72 °C for 10 min. A total of 16 PCR reactions were performed and pooled for each DNA template. The ~450 bp PCR products were purified, digested with *Sfi*I restriction enzyme, and purified again. The inserts were then ligated into pMED1 phagemid vector (Arbabi-Ghahroudi *et al*. 2009a) which had also been digested with *Sfi*I restriction enzyme (with the addition of *Xho*I and *Pst*I restriction enzymes to reduce vector self-ligation) and the ligated vectors purified.

Finally, 20 × 50 μL aliquots of TG1 *E. coli* (Agilent Technologies; Santa Clara, CA, USA; Cat. #: 200123) were each transformed with 5 μL (~350 ng) ligated pMED1 using a BioRad® Gene Pulser electroporation device, titered and grown overnight at 37 °C, 250 rpm in 2xYT supplemented with 100 μg/mL ampicillin and 2% (w/v) glucose, as described by Hussack *et al*. (2012a) and Baral *et al*. (2013). The following morning, the bacteria were pelleted, resuspended in 20 mL 2xYT supplemented with 100 μg/mL ampicillin and 2% (w/v) glucose + 20 mL 50% (v/v) glycerol, and stored in 1 mL aliquots at −80 °C. Colonies on the titer plates were counted as well as analyzed using colony-PCR (15 μL reactions consisting of 1.5 pmol −96gIII primer, 1.5 pmol M13RP primer, 0.4 μL 10 mM dNTPs, 0.15μL GenScript *Taq* DNA polymerase (GenScript; Cat. #: E00007), 1.5 μL Taq kit 10x buffer, 12.65 μL nuclease free H2O, and a single colony pick using thermocycler parameters of 94 °C for 5 min, 35 cycles of 95 °C for 30 s, 55 °C for 30 s and 72 °C for 45 s, and a final extension of 72 °C for 10 min), to determine the total and functional library sizes. Two aliquots of library cells were thawed, infected with ~7.2 × 10^11^ plaque-forming units M13KO7 helper phage (New England BioLabs; Ipswich, MA, USA; Cat. #: N0315S), grown overnight, and the phage particles harvested from the culture supernatant and titered, as described by Hussack *et al*. (2011), Hussack *et al*. (2012a), and Baral *et al*. (2013), to create the input library phage for panning. Aliquots of ~100-200 μL input phage in PBS were kept at −80 °C for long-term storage.

### 2.5 Subtractive panning

The screening of both VHH phage display libraries for binders to the recombinant HPV16 E6 proteins was completed using panning techniques (Schmitz *et al*. 2000) routinely employed by us and others (Arbabi-Ghahroudi *et al*. 1997, Hussack *et al*. 2011, Hussack *et al*. 2012a, Hussack *et al*. 2012b, Baral *et al*. 2013, Pardon *et al*. 2014). In the first round, two Immuno™ Breakable Module MaxiSorp™ wells (Thermo Fisher Scientific; Cat. #: 12-565-134) were coated with 10 μg of MBP, one was coated with a mix of 5 μg His6MBP-4C/4S E6 + 5 μg His6MBP-F47R 4C/4S E6, and one was coated with PBS (100 μL/well; antigens diluted in PBS) overnight at 4 °C. All wells were then blocked with 2% milk powder in PBS for 1 h at 37 °C. To subtract the MBP binders, 100 μL Library #1 input phage (~1 × 10^12^ phage) + 100 μL 4% milk powder in PBS were added to each of the two blocked, MBP-coated wells and incubated for 1 h at room temperature. Phage from one MBP-coated well were then transferred to the blocked, His6MBP-E6-coated well and phage from the other MBP-coated well were transferred to the blocked, “PBS-coated” well. Phage were incubated for 30 min at room temperature. The same procedure was completed in parallel using Library #2 input phage. Following washing with PBS-T, bound phage were collected in two sequential elutions: 1) 100 mM triethylamine neutralized with 1 M Tris-HCl pH 7.4 and 2) 100 mM glycine pH 2.0 neutralized with 2 M Tris base, and both elutions pooled. Exponentially growing TG1 cells were infected with the eluted phage and an aliquot taken for titration. The remaining infected cells were then superinfected with ~20x excess M13KO7 helper phage, grown overnight at 32 °C, and the amplified phage purified, as described by Hussack *et al*. (2011), Hussack *et al*. (2012a), and Baral *et al*. (2013), to create input phage for the second round of panning. Here, amplified phage were quantified spectrophotometrically using the Antibody Design™ Laboratories Phage Concentration Calculator (http://www.abdesignlabs.com/technical-resources/phage-calculator/) which employs a formula based on the measurements by Day and Wiseman (1978):

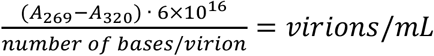

A second round of panning was completed as described above, except that each subtraction well was coated with an increased amount (20 μg) of MBP, each target antigen well was coated with a decreased amount (2.5 μg His6MBP-4C/4S E6 + 2.5 μg His6MBP-F47R 4C/4S E6) of E6 proteins, and the blocking buffer switched to SuperBlock™ (Thermo Fisher Scientific; Cat. #: PI37580), to increase selective pressure for higher affinity binders and prevent enrichment of VHHs with affinity for milk powder or MBP.

### 2.6 Phage ELISA

To examine progress after each round of panning, 8 randomly-selected colonies from the eluted phage titer plates for each library were first analyzed using colony-PCR (as described in Section 2.4) to confirm the presence of VHH inserts. Next, 48 colonies from the round 1 eluted phage titer plates or 96 colonies from the round 2 eluted phage titer plates for each library were sequenced using the M13RP primer. VHH clones of interest were further characterized for their ability to bind recombinant HPV16 E6 using phage ELISA. As similarly described by Hussack *et al*. (2012a) and Baral *et al*. (2013), 1 μg of His6MBP-4C/4S E6, His6MBP-F47R 4C/4S E6, or MBP (100 μL/well; antigens diluted in PBS) or 100 μL/well PBS alone was coated in Nunc™ MaxiSorp™ 96-well plates overnight at 4 °C. The wells were blocked with 5% milk powder in PBS-T at 37 °C, prior to incubation with each respective VHH-displaying phage amplified from the colonies sequenced above for 1 h at 37 °C. The wells were then washed with PBS-T and incubated with a mouse anti-M13 + HRP antibody (GE Healthcare; Mississauga, ON, Canada; Cat. #: 27-9421-01) diluted 1:5 000 in PBS for 1 h at room temperature. The wells were again washed with PBS-T and then incubated with TMB as described in Section 2.3. All clones of interest were tested under these conditions in two separate experiments.

### 2.7 VHH expression and purification

DNA inserts consisting of an OmpA leader (to direct VHH expression to the periplasm of *E. coli* (Movva *et al*. 1980)) + VHH + human influenza hemagglutinin (HA)-His6 tags were synthesized for 26 VHHs which yielded notable E6-specific phage ELISA signals as well as for one VHH which demonstrated MBP-binding, and these inserts were subcloned into the pSJF2H expression vector (Arbabi-Ghahroudi *et al*. 2009b) via the EcoRI and *Bam*HI restriction sites through services available at GenScript. Aliquots of 5 μL Zymo Mix and Go!™ chemically competent TG1 *E. coli* (Cedarlane Laboratories; Cat. #: T3017) were transformed with ~50 ng insert-containing pSJF2H vector, according to supplier instructions. The transformed cells were grown in 250 mL 2xYT media supplemented with 100 μg/mL ampicillin and 0.1% (w/v) glucose and VHH expression induced with 0.8 mM IPTG overnight at 37 °C. Periplasmic extraction using TES buffer, IMAC purification, and calculation of VHH concentration were completed as similarly described by Hussack *et al*. (2011), except both incubations on ice during TES extraction were increased from 30 min to 1 h. The purified VHHs were then aliquoted and stored at −20 °C long-term, following their analysis with reducing SDS-PAGE.

### 2.8 E6-detection capability of purified VHHs

To characterize the ability of the purified VHHs to detect recombinant HPV16 E6, ELISA, reducing SDS-PAGE and native PAGE Western blots, as well as dot blots were performed. For the ELISA, 0.5 μg of His6MBP-4C/4S E6, His6MBP-F47R 4C/4S E6, or MBP (100 μL/well; His6MBP-E6 antigens diluted in their storage buffer, MBP diluted in PBS) or 100 μL/well His6MBP-E6 protein storage buffer alone was coated in Nunc™ MaxiSorp™ 96-well plates overnight at 4 °C. The wells were blocked with 5% milk powder in PBS-T at 37 °C, prior to application of 100 μg/mL of each VHH diluted in PBS (100 μL/well) for 1 h at room temperature. The wells were then washed with PBS-T and incubated with a rabbit anti-His6 + HRP antibody (Cedarlane Laboratories; Cat. #: A-190-114P) diluted 1:20 000 in PBS (100 μL/well) for 1 h at room temperature. The wells were again washed with PBS-T and then incubated with TMB as described in Section 2.3.

For Western blots run under reducing conditions, 10 ng/lane of His6MBP-4C/4S E6, His6MBP-F47R 4C/4S E6, or MBP as well as 5 ng/lane HAHis6-tagged VHH (as a positive secondary antibody control) were separated on Mini-PROTEAN® TGX™ precast 4-20% gradient polyacrylamide gels (BioRad; Mississauga, ON, Canada; Cat. #: 4561096) for ~70 min at 120 V and transferred to PVDF membrane for 1 h at 100 V. The membranes were blocked with 5% milk powder in TBS-T for 1 h at room temperature followed by overnight 4 °C incubation with 2.7 μg/mL purified VHH or mouse monoclonal anti-HPV16 E6 antibody (clone 6F4; kind gift from Arbor Vita Corporation; Fremont, CA, USA) diluted in blocking buffer. As this concentration of mAb yields reproducible detection from HPV16-positive cell lysates in our lab (Togtema *et al*. 2018), it was employed as our initial test concentration here. The membranes were washed with TBS-T, followed by incubation with a goat polyclonal anti-HA tag + HRP secondary antibody (Abcam; Toronto, ON, Canada; Cat. #: ab1265) diluted 1:5 000, a mouse monoclonal anti-HA tag + HRP secondary antibody (Thermo Fisher Scientific; Cat. #: 26183-HRP) diluted 1: 1 000-1:2 000 (depending on individual lot characteristics), or a goat anti-mouse IgG + HRP secondary antibody diluted 1:2 000 in blocking buffer for 1 h at room temperature, as appropriate. The membranes were again washed in TBS-T and chemiluminescence detection completed as described in Togtema *et al*. (2018). All VHHs were tested under these conditions in a minimum of two separate experiments.

For the native PAGE Western blots, 5 μg/lane of His6MBP-4C/4S E6, His6MBP-F47R 4C/4S E6, or MBP as well as 2.5 μg/lane HA-His6-tagged VHH (as a positive secondary antibody control) were separated on 10% polyacrylamide gels without SDS for 2.5 h at 100 V, 4 °C and transferred (in buffer without MeOH) to PVDF membrane for ~16 h at 20 V, 4 °C. Immunoblotting was then performed as described above. All VHHs were tested under these conditions in two separate experiments.

For the dot blots, 2 μg of His6MBP-4C/4S E6, His6MBP-F47R 4C/4S E6, or MBP was spotted on a nitrocellulose membrane and allowed to dry for 1 h at room temperature. The membrane was then blocked with 5% milk powder in PBS-T (0.01%) for 1 h at room temperature followed by overnight 4 °C incubation with 2.7 - 10.8 μg/mL purified VHH diluted in PBS-T. The membranes were washed with PBS-T, followed by incubation with an anti-HA tag + HRP secondary antibody as described above. The membranes were again washed in PBS-T and chemiluminescence detection completed as indicated above.

### 2.9 Size exclusion chromatography and surface plasmon resonance analyses

Prior to surface plasmon resonance (SPR) experiments, the VHHs and His6MBP-E6 proteins were size exclusion chromatography (SEC) purified. The VHHs (300-500 μg) were purified using a Superdex 75 Increase 10/300 GL column (GE Healthcare) while the His6MBP-E6 proteins (300 μg) were purified using a Superdex 200 Increase 10/300 GL column (GE Healthcare). A flow rate of 0.5 mL/min and HBS-EP+ buffer (10 mM HEPES pH 7.4, 150 mM NaCl, 3 mM EDTA, 0. 5% (v/v) P20 surfactant) were used for all experiments. Fractions of 0.5 mL volume were collected and the concentration determined through absorbance measurements at 280 nm.

All SPR analyses were performed on a Biacore T200 instrument (GE Healthcare) at 25 °C, using HBS-EP+ as running buffer and Series S CM5 sensor chips (GE Healthcare). All data were reference-flow cell subtracted and analyzed using Biacore T200 software v3.0 (GE Healthcare). Two SPR assay formats were performed. In the first assay, E6 proteins (His6MBP-4C/4S E6, His6MBP-F47R 4C/4S E6) and a control MBP protein (His6MBP-intimin) were coupled to a CM5 chip through standard amine coupling. Briefly, ~1200 RUs of each protein were immobilized in 10 mM acetate buffer pH 5.0 (His6MBP-E6 proteins) or pH 4.5 (His6MBP-intimin), producing surfaces with theoretical *R*_max_s ranging from 260-300 RUs for the VHHs. VHHs (A01, A05, A09, A46, 2A12, 2A17, C26) were injected at a single concentration of 500 nM and an HPV16 E6-specific mAb (clone 6F4; as described in Section 2.8) was injected at 100 nM, all at a flow rate of 50 μL/min for 180 s followed by 600 s of dissociation. Surfaces were regenerated with a 120 s pulse of 10 mM glycine pH 1.5 at a flow rate of 30 μL/min. In the second assay, VHHs (A01, A05, A09, 2A12, 2A17, C26) and the HPV16 E6-specific mAb were coupled to Series S CM5 sensor chips using similar conditions reported above (10 mM acetate buffer pH 5.0). Approximately 400 RUs of each VHH or mAb were immobilized. Non-SEC purified His6MBP-E6 or His6MBP-intimin was injected at a single concentration of 500 nM (over VHH surfaces) or 100 nM (over the mAb surface) to determine surface activity. Next, using single-cycle kinetics, a concentration range (500 nM-31.5 nM) of SEC-purified His6MBP-4C/4S E6 or His6MBP-F47R 4C/4S E6 was injected over each surface at a flow rate of 40 μL/min for 180 s followed by 600 s of dissociation to determine affinities and kinetics. Sensorgrams were reference subtracted and fit to a 1:1 binding model. Surface regeneration conditions were identical to the first assay described above.

## 3. Results

### 3.1 A specific heavy-chain IgG response was induced following each llama immunization

To maximize our odds of isolating high-affinity VHHs targeting HPV16 E6, we performed two separate llama immunizations with different recombinant antigens. The first llama (Immunization #1) was injected with His6-GenScript E6: a variant of HPV16 E6 corresponding to that found in the cervical carcinoma-derived HPV16-positive cell line CaSki (Pattillo *et al*. 1977, Meissner *et al*. 1999, Zehbe *et al*. 2009) with an N-terminal His6 tag (Figure 3). The second llama (Immunization #2) was injected with a mixture of His6MBP-4C/4S E6 and His6MBP-F47R 4C/4S E6, which are solubility-enhanced mutants of HPV16 E6 each with an N-terminal His6-MBP tag followed by a TEV protease cleavage site (Zanier *et al*. 2012, Zanier *et al*. 2013, Ramirez *et al*. 2015, Martinez-Zapien *et al*. 2016) (Figure 3). These immunization approaches were intended to be complementary, as the first used a variant of E6 which was naturally-occurring but minimally-soluble in recombinant form, and had only a small His6 tag added; whereas the second used lab-engineered mutants of E6 which demonstrated improved recombinant solubility, but also had a large (MBP; ~42 kDa) fusion add-on as well as the His6 tag.

Induction of the desired antibody responses was initially assessed by screening the total llama serum by ELISA. As anticipated, serum collected 35 and 42 days following the initial immunization contained increased amounts of IgG which reacted with the injected recombinant antigen(s) compared to pre-immune serum collected on Day 0, for both Immunization #1 (Figure 4A) and Immunization #2 (Figure 4B). The pre-immune serum from Immunization #2 also contained some IgG with minor reactivity for the His6MBP-E6 proteins. This may perhaps be the result of a previous immune encounter the animal had with the E6 protein of a distantly-related llama papillomavirus (Schulman *et al*. 2003, Ure *et al*. 2011) or, more likely, with the bacterial MBP (Boos and Shuman 1998).

**Figure 4.**
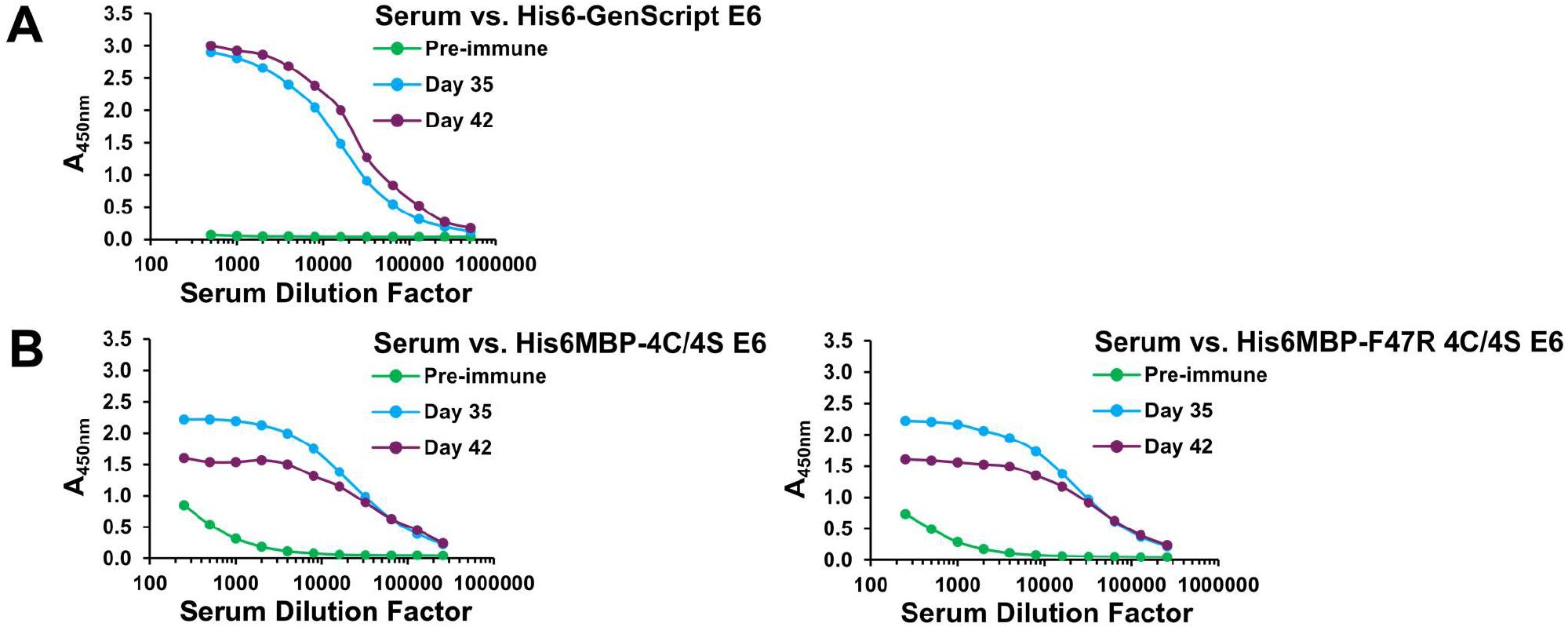
Total serum response following each llama immunization. Following both Immunization #1 **(A)** and Immunization #2 **(B)**, ELISAs demonstrated that serum collected on Days 35 and 42 contained an increased amount of IgG which reacted with the injected recombinant antigen(s) compared to pre-immune serum collected on Day 0, indicating specific immune responses had been induced.

Next, Day 0 and Day 42 total serum from both immunizations was fractionated using protein A and protein G affinity chromatography (Hussack *et al*. 2012a, Baral *et al*. 2013) (Figure 5A). For simplicity, Day 35 serum was omitted from this step as its total response was similar to that of Day 42 serum in both instances. Subsequent ELISAs confirmed that, in addition to conventional IgG, heavy-chain IgG were also involved in the observed positive total serum responses for both Immunization #1 (Figure 5B) and Immunization #2 (Figure 5C). In particular, for Immunization #1, only the Day 42 heavy-chain IgG3 (G1) fraction displayed a notable increase in reactivity for the injected recombinant antigen compared to its corresponding Day 0 fraction. Whereas, for Immunization #2, an increase was observed in all three heavy-chain IgG3 (G1), IgG2a (A1), and IgG2b/c (A2) fractions. Different amounts of contaminating IgM in the Day 42 versus Day 0 A2 serum fractions, which co-elutes during the chromatography process, may have skewed our ability to detect a positive heavy-chain IgG2b/c response in the case of Immunization #1. However, generally weak heavy-chain IgG2a/b/c immune responses have also been observed by us previously and have not affected our ability to isolate high affinity VHHs from the resulting phage display libraries.

**Figure 5.**
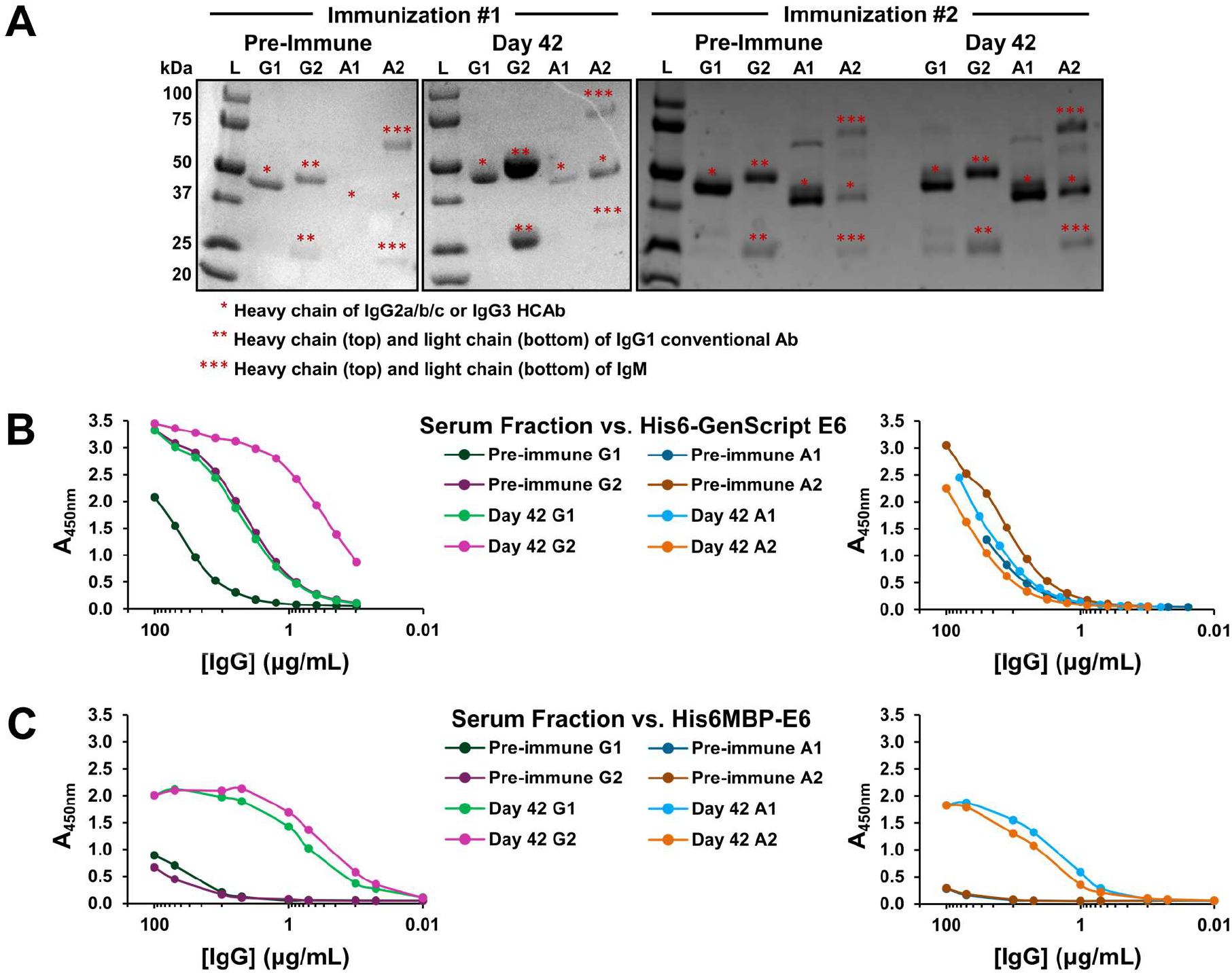
Serum fractionation and confirmation of a heavy-chain IgG (HCAb) immune response following each llama immunization. **(A)** Reducing SDS-PAGE was used to analyze the purity of the pre-immune and Day 42 serum fractions: G1 fraction = IgG3 HCAb, G2 fraction = IgG1 conventional antibody, A1 fraction = IgG2a HCAb, and A2 fraction = IgG2b/c HCAb. Approximately 2.0-2.5 μg of sample was loaded in each lane, with the exception of some of the more dilute A1 and A2 fractions for which smaller, but still visualizable, amounts were loaded. IgM which sometimes co-elutes with IgG2b/c was noted in all A2 fractions examined here. In the Immunization #2 pre-immune and Day 42 A1 serum fractions, an additional band is visible below the 75 kDa marker which may consist of residual unreduced HCAb. **(B)** ELISAs demonstrated that Immunization #1 Day 42 G1 and G2 serum fractions showed increased recognition for the injected recombinant antigen compared to the corresponding pre-immune serum fractions. An increased response was also observed for all four Immunization #2 Day 42 serum fractions **(C)**, confirming HCAb immune responses had been induced in both llamas.

### 3.2 VHH phage display libraries corresponding to each llama immunization were successfully constructed and enriched for antigen-specific binders using subtractive panning

Following confirmation of heavy-chain IgG responses, a VHH phage display library corresponding to each llama immunization (i.e., Library #1: His6-GenScript E6 immunization and Library #2: His6MBP-E6 immunization) was constructed from frozen lymphocytes collected on Day 42. Using well-established techniques (Hussack *et al*. 2011, Hussack *et al*. 2012a, Baral *et al*. 2013), VHH DNA was PCR amplified, subcloned into the pMED1 phagemid vector (Arbabi-Ghahroudi *et al*. 2009a), and transformed into TG1 *E. coli*. It was estimated that Library #1 had a functional size of ~3.4 × 10^8^ transformants (as calculated by 3.8 × 10^8^ total transformants multiplied by the 90.6% of colonies determined to contain VHH inserts) and Library #2 had a functional size of ~1.7 × 10^7^ transformants (as calculated by 1.8 × 10^7^ total transformants multiplied by the 93.8% of colonies determined to contain VHH inserts).

As the successful usage of recombinant HPV16 E6 in molecular biology applications was notably facilitated by the solubility-enhanced mutants (Suarez and Travé 2018), we chose to exclude 6His-GenScript E6 from use as a target antigen during the subsequent panning and VHH characterization assays. Accordingly, each library was then individually panned against a mixture of both His6MBP-E6 proteins. To select against MBP-specific VHHs, a subtractive panning technique (Hussack *et al*. 2012b) was employed whereby the input phage were first incubated in a well coated with MBP before being transferred to His6MBP-E6 or PBS coated wells. Two rounds of panning were completed for each library, with the amount of protein in the MBP-coated subtractive well increased and the amount of protein in the His6MBP-E6-coated well decreased during the second round, to maximize selective pressure for high affinity, E6-specific VHHs. The blocking reagent was also switched during the second round, to prevent unintentional enrichment for VHHs with affinity to it. The number of VHH-displaying phages eluted from the His6MBP-E6 coated wells during each round of panning are reported in Table 1. For Library #2, they were ~157.1x and ~166666.7x greater than the number eluted from corresponding control wells coated with PBS to measure the background number of VHH-displaying phage with affinity for the plastic or blocking reagent. For Library #1, they were only ~12.5x and ~1.2x greater. These smaller numbers may reflect the fact that Library #1 was panned against the His6MBP-E6 proteins but was generated by immunization with His6-GenScript E6, whereas Library #2 was both panned against and generated by immunization with the His6MBP-E6 proteins.

**Table 1.**
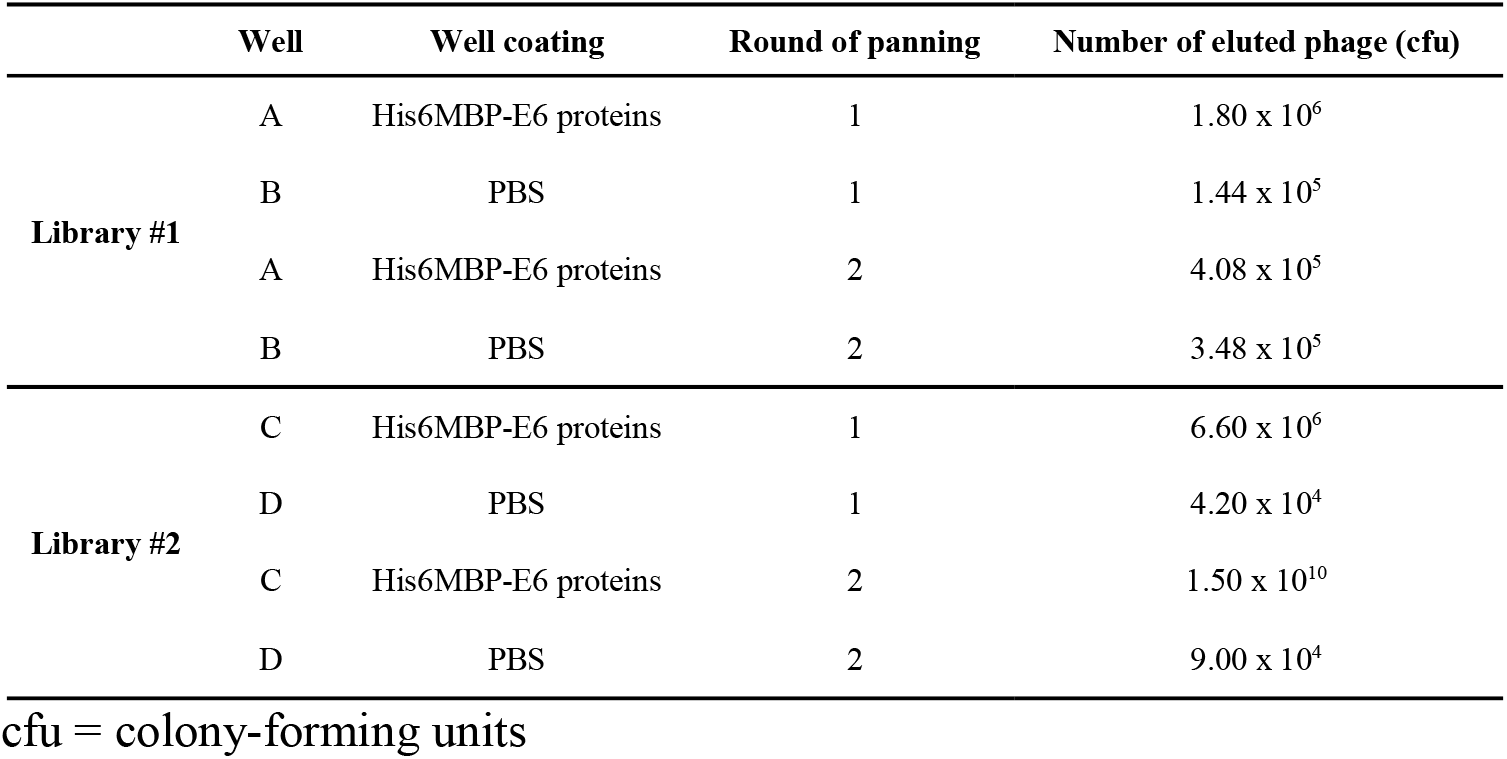
Number of VHH-displaying phage eluted during each round of panning. cfu = colony-forming units

|  | Well | Well coating | Round of panning | Number of eluted phage (cfu) |
| --- | --- | --- | --- | --- |
| **Library #1** | A | His6MBP-E6 proteins | 1 | $1.80 \times 10^6$ |
| | B | PBS | 1 | $1.44 \times 10^5$ |
| | A | His6MBP-E6 proteins | 2 | $4.08 \times 10^5$ |
| | B | PBS | 2 | $3.48 \times 10^5$ |
| **Library #2** | C | His6MBP-E6 proteins | 1 | $6.60 \times 10^6$ |
| | D | PBS | 1 | $4.20 \times 10^4$ |
| | C | His6MBP-E6 proteins | 2 | $1.50 \times 10^{10}$ |
| | D | PBS | 2 | $9.00 \times 10^4$ |

cfu = colony-forming units

### 3.3 The eluted VHH-displaying phage were further characterized using phage ELISA

Following panning, the VHH phagemid inserts of 48 random colonies from the round 1 eluted phage titer plates and of 96 random colonies from the round 2 eluted phage titer plates for each library were arbitrarily numbered and sequenced. As indicated in Table 1, clones beginning with “A” or “C” were isolated from Library #1 or Library #2 round 1 eluted phage titer plates, respectively, and clones beginning with “2A” or “2C” were isolated from Library #1 or Library #2 round 2 eluted phage titer plates, respectively. VHH-displaying phage were then amplified for more than 40 clones with different complementarity-determining region 3 (CDR3) sequences (the main region of the VHH involved in interaction with the antigen (Steeland *et al*. 2016)), and these VHHs were further characterized for their ability to bind to the His6MBP-E6 proteins by phage ELISA (Hussack *et al*. 2012a, Baral *et al*. 2013).

Of those investigated, 26 clones which generated a strong signal in wells coated with either His6MBP-E6 protein but not in wells coated with MBP or PBS were considered to be potential E6 binders and were selected for soluble expression and purification (Figure 6). The occurrences of additional clones with the same CDR3 sequences as these VHHs are detailed in Supplementary Table 1. Nine of our 26 candidates had CDR3 sequences which were identified multiple times and three (A47, C11, and C38) had CDR3 sequences which were identified in both libraries. Despite the implementation of subtractive panning techniques, the majority of clones identified from the Library #2, round 1 and round 2 eluted phage titer plates were MBP binders which generated signal in wells coated with MBP, in addition to wells coated with either His6MBP-E6 protein. One of these clones, C26, was also selected for use as an MBP-binding control VHH in the forthcoming assays (Supplementary Figure 1A).

**Figure 6.**
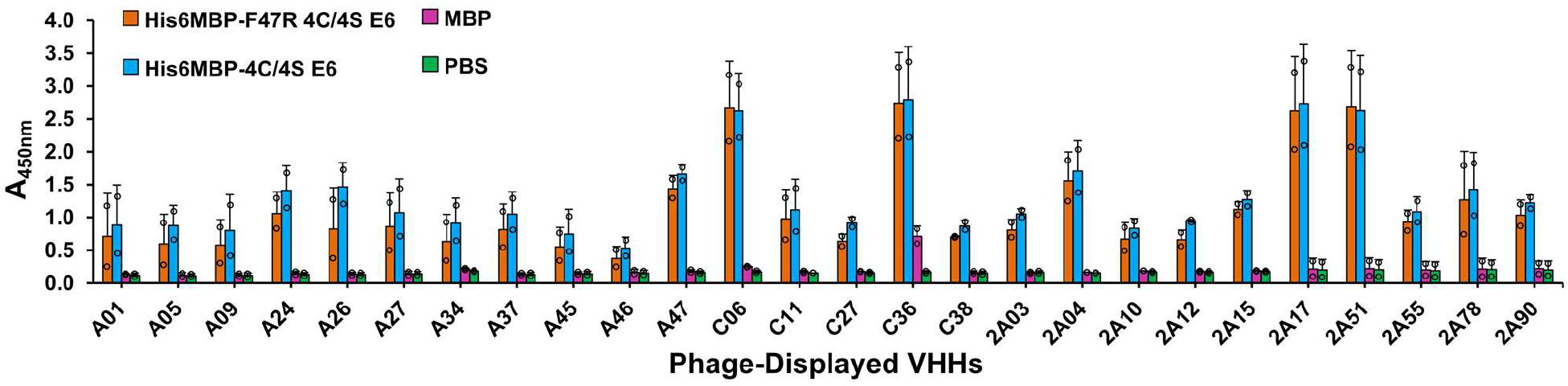
Phage ELISA. Clones of interest from the eluted phage titer plates for both libraries were considered to be potential E6 binders if a strong signal was observed following incubation of phage displaying that VHH in wells coated with either His6MBP-E6 protein but not in wells coated with MBP or PBS. Twenty-six unique clones exhibiting this phage ELISA profile were chosen for soluble expression and purification. Clones beginning with “A” or “C” were isolated from Library #1 or Library #2 round 1 eluted phage titer plates, respectively. Clones beginning with “2A” were isolated from Library #1 round 2 eluted phage titer plates. Data represent mean + standard deviation, with individual data points overlaid (n = 2).

### 3.4 Expression of VHHs

The 26 potential E6-binding clones identified by phage ELISA as well as the one MBP-binding clone (C26) were next expressed as soluble VHHs with a C-terminal HA-His6 tag, following subcloning of each corresponding DNA insert into the pSJF2H vector (Arbabi-Ghahroudi *et al*. 2009b). VHH expression was directed to the *E. coli* periplasm using an OmpA leader (Movva *et al*. 1980), which allowed for proper disulfide bond formation (De Marco 2009) as well as ease of extraction through osmotic shock rather than total cell lysis (Baral *et al*. 2013), and the VHHs were purified from the resulting extract using IMAC (Hussack *et al*. 2011) (Figure 7A and Supplementary Figure 1B). Five of the 26 potential E6-binding VHHs (clones A26, C06, C27, 2A55, and 2A90) did not express with our overnight approach. The expression of clone A24 was also quite low relative to the other VHHs. Hence, due to limited starting material, these six clones were excluded from further analyses.

**Figure 7.**
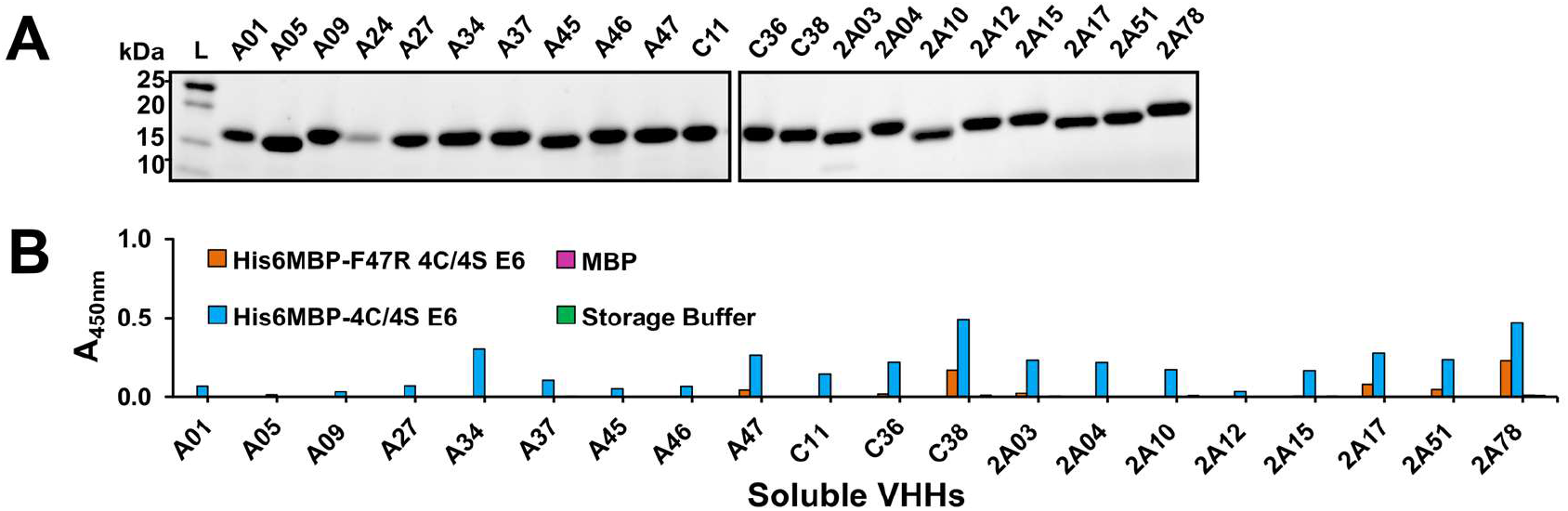
VHHpurification and determination of their ability to detect recombinant HPV16 E6. **(A)** Reducing SDS-PAGE was used to analyze the purity of the VHHs following IMAC purification. Approximately 3 μg of sample was loaded in each lane, with the exception of the more dilute clones A01, A24, C38, and 2A17 for which amounts ranging from approximately 1-2.5 μg were loaded. **(B)** An ELISA using 100 μg/mL of each clone demonstrated that the purified VHHs were functional but varied in their ability to detect the His6MBP-E6proteins.

To initially gauge the ability of the purified VHHs to detect the His6MBP-E6 proteins, we performed an ELISA using 100 μg/mL of each clone as the primary antibody (Figure 7B). Due to the presence of a His6 tag on our antigens as well as our VHHs, we first tried an anti-HA + HRP secondary antibody. This, however, resulted in very weak signal with high non-specific background. Instead, we then used an anti-His6 + HRP secondary antibody. The background signal originating from the N-terminal His6 tags of the recombinant His6MBP-E6 proteins was accounted for by subtracting the absorbance of His6MBP-E6, MBP or buffer-coated control wells, which were incubated with PBS instead of VHH, from the corresponding wells incubated with both a VHH and secondary antibody. The clones demonstrated varying levels of binding to both His6MBP-E6 proteins but, overall, the signal was notably weaker than that obtained by phage ELISA. This may be caused by less amplification of the signal from bound VHH by the anti-His6 + HRP secondary antibody relative to that generated by the anti-M13 + HRP secondary antibody, which targets a coat protein present in multiple copies per phage (Baral *et al*. 2013). It may also, in part, be an artefact of our subtractive method of analysis in this instance. As expected, C26 bound to MBP, in addition to both His6MBP-E6 proteins (Supplementary Figure 1C).

### 3.5 Characterization of VHHs using Western blotting under denaturing and native conditions

To narrow down our current pool of E6-binding clones to those we first sought to analyze using SPR, each VHH was tested in both reducing SDS-PAGE and native PAGE Western blots. This allowed us to further characterize their interactions with the recombinant His6MBP-E6 proteins in two additional, widely-used molecular biology assays and provided insight into the ability of each VHH to detect a linear or conformational epitope (as similarly described by Hussack *et al*. (2011) and Hussack *et al*. (2012a)). Both blot types were repeated at least twice per clone, using purified VHH as the primary antibody followed by application of an anti-HA tag + HRP secondary antibody. A sample of purified VHH was also run on each gel, to rule out inconsistent functionality of the secondary antibody as a cause for any negative results. As a positive control, the HPV16 E6-specific 6F4 mAb (Giovane *et al*. 1999) was also tested in these same assays, together with an anti-mouse + HRP secondary antibody.

In their current monomeric format, none of the potential E6-binding VHHs yielded a detectable signal under denaturing conditions. However, under native conditions, clone 2A17 yielded reproducible bands in the lanes loaded with either His6MBP-E6 protein but not in the lane loaded with MBP, indicating that it interacts with a conformational epitope on the E6 portion of the antigen. Comparatively, we found the 6F4 mAb bound to both denatured and native His6MBP-E6 proteins, indicating that it interacts with a linear E6 epitope (Hussack *et al*. 2012), as was originally reported by Giovane *et al*. (1999) and Choulier *et al*. (2002). Clone C26 yielded reproducible bands in lanes loaded with MBP, in addition to lanes loaded with either His6MBP-E6 protein, confirming its status as an MBP-binding VHH. It too appears to interact with a linear epitope, as indicated by signal in both types of blots. As expected, none of the secondary antibodies bound to our recombinant antigens in the absence of primary antibody. These results are summarized in Figure 8.

**Figure 8.**
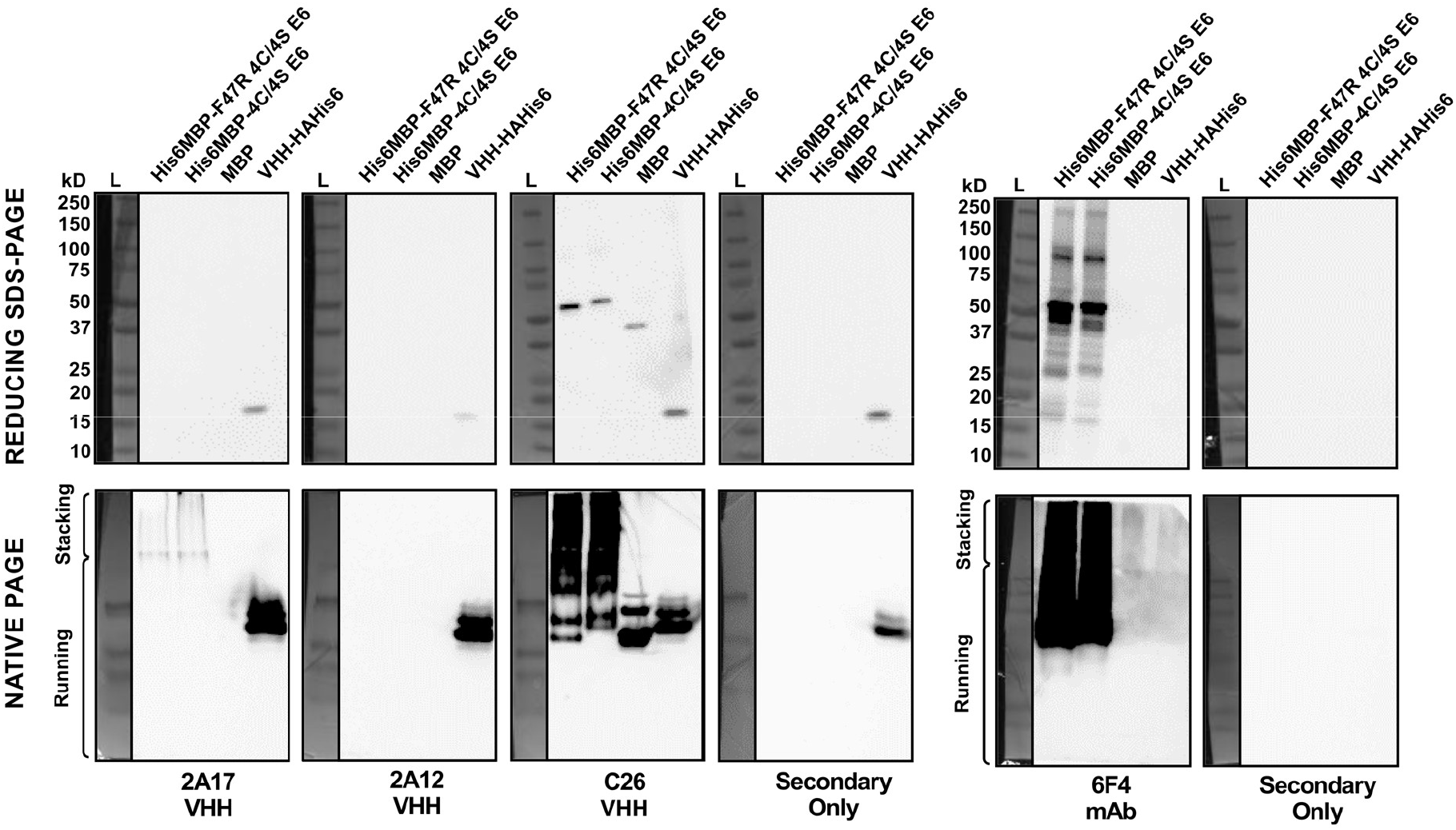
Reducing SDS-PAGE and native PAGE Western blots. When considered altogether, the assay results indicated clone 2A17 interacts with a conformational epitope on the E6 portion of the His6MBP-E6 proteins; whereas none of the other purified VHHs yielded detectable signal (clone 2A12 shown as an example). In contrast, the HPV16 E6-specifiic 6F4 mouse mAb interacts with a linear E6 epitope. Clone C26 was also further confirmed as an MBP binder which likely also interacts with a linear epitope. For reducing SDS-PAGE, 10 ng/lane of His6MBP-F47R 4C/4S E6, His6MBP-4C/4S E6, and MBP as well as 5 ng/lane of purified VHH 2A04 was loaded. For native PAGE, 5 μg/lane of His6MBP-F47R 4C/4S E6, His6MBP-4C/4S E6, and MBP as well as 2.5 μg/lane of purified VHH A46 was loaded. Note, for the native PAGE Western blots, a ladder was used only as an indicator of protein transfer and does not reflect the molecular weight of the resolved proteins.

### 3.6 Surface plasmon resonance analyses demonstrated several VHHs bound recombinant E6 with nanomolar affinity

Next, we sought to determine with what affinity clone 2A17 bound to the recombinant E6 proteins. Based on the above described Western blot results, we also included clones A01, A05, A09, and 2A12 as expected negative controls, the HPV16 E6-specific 6F4 mAb as a positive control, as well as clone C26 as an MBP-binding control. The initial assay setup involved amine coupling of both His6MBP-E6 proteins as well as a control His6MBP-tagged intimin protein to the SPR sensor chip. SEC-purified VHHs (all were SEC purified except A09, due to its low concentration and yield) or the 6F4 mAb (not SEC purified) were then flowed over the chip to assess “Yes/No” binding (data not shown). No notable binding to any of the recombinant proteins by VHHs A01, A05, A09, 2A12, or 2A17 was detected (data not shown). However, the 6F4 mAb bound to both His6MBP-E6 proteins and the C26 VHH bound to all three recombinant proteins containing MBP as expected (Figure 9A).

**Figure 9.**
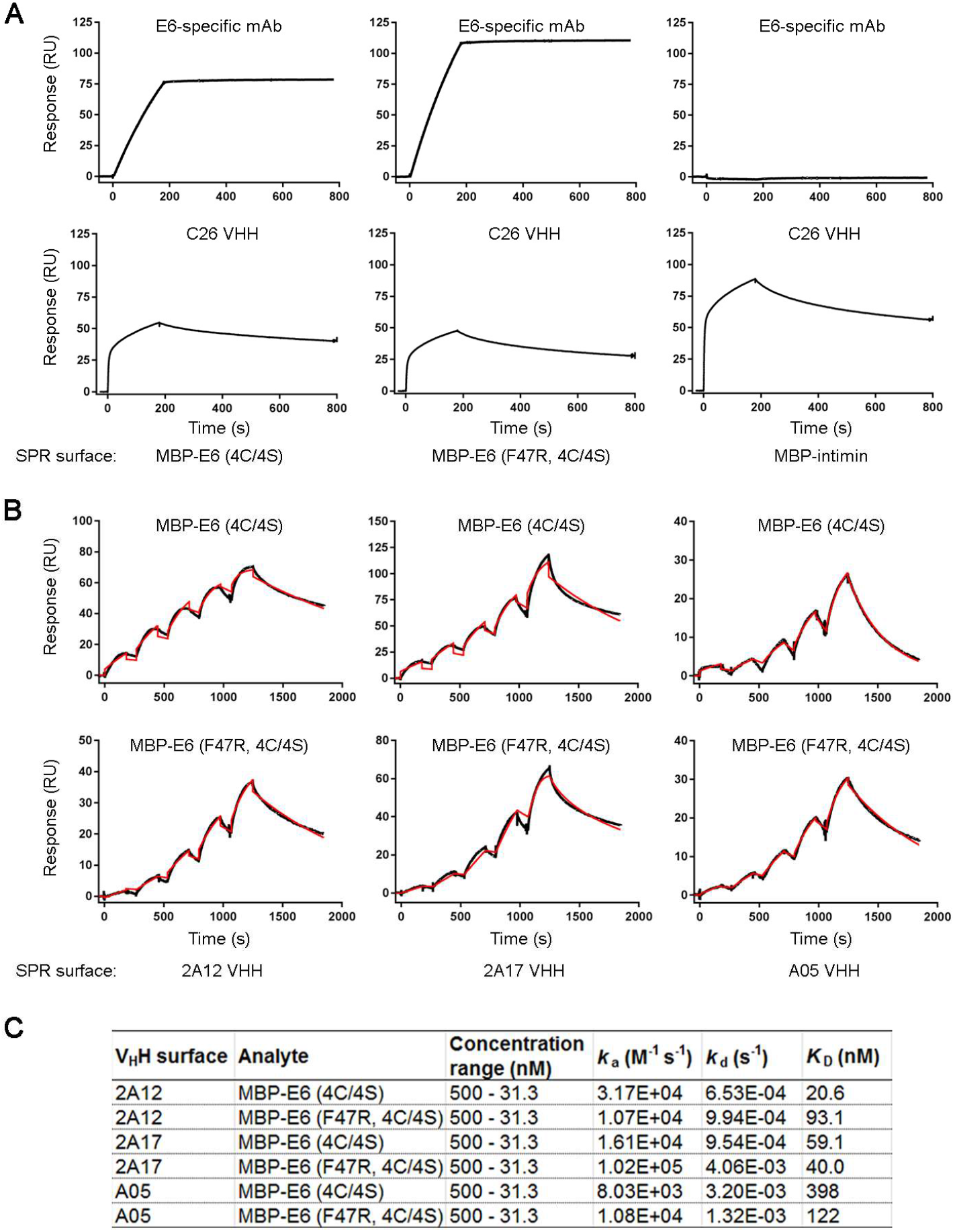
Surface plasmon resonance (SPR) binding assays demonstrating the interaction of the His6MBP-E6 proteins and VHHs. **(A)** SPR surfaces with His6MBP-E6 or control MBP (His6MBP-intimin) were created and VHHs or mAbs flowed over. The HPV16 E6-specific 6F4 mAb bound E6-containing surfaces and not the MBP control surface while the C26 VHH bound all MBP-containing surfaces. All other VHHs tested did not bind any surface (data not shown). **(B)** VHHs were immobilized and SEC-purified E6 proteins flowed over using single-cycle kinetics to determine affinities and kinetics. Black lines represent raw data and red lines represent 1:1 binding model fits. **(C)** Summary of VHH binding affinities calculated from the sensorgrams in **(B)**.

To rule out the possibility that amine coupling was impacting the integrity of the E6 proteins, a second SPR assay was performed where the VHHs or 6F4 mAb were amine-coupled to the SPR sensor chip and His6MBP-E6 proteins flowed over the chip. Using this assay format, E6-specific binding was observed for VHHs A05, 2A12, and 2A17 (they bound to both His6MBP-E6 proteins but not the control His6MBP-tagged intimin protein) (data not shown). Clones A01 and A09 did not bind to any of the recombinant proteins (data not shown). As expected, the 6F4 mAb bound to both His6MBP-E6 proteins and clone C26 bound to all three recombinant proteins (data not shown). We hypothesized that the harsh coupling conditions in the initial assay setup may have disrupted the conformational epitopes detected by these VHHs but did not affect the binding of the 6F4 mAb, which, in contrast, detects a linear epitope at the N-terminus of the E6 protein (Giovane *et al*. 1999, Choulier *et al*. 2002). Using this second assay orientation, single-cycle kinetics data were then collected for the VHHs to determine affinities and kinetics. Confirming and elaborating upon the “Yes/No” binding results, we found clones A05, 2A12, and 2A17 bound both His6MBP-E6 proteins with nanomolar range affinities (Figure 9B and C) but did not bind the control His6MBP-tagged intimin protein.

As we were able to detect VHHs A05 and 2A12 binding the recombinant E6 proteins using SPR (a label-free assay) but not with our above described Western blots under native conditions (which rely on the VHH HA tags), we first confirmed the HA tags were still intact on our stored stocks of purified VHHs by resolving samples of each clone by reducing SDS-PAGE, transferring them to PVDF and then probing the membranes with an anti-HA + HRP antibody (Supplementary Figure 2). Next, the recombinant His6MBP-E6 proteins as well as the recombinant MBP protein were spotted onto nitrocellulose membranes in dot blot format and increased VHH concentrations were tested. We were able to obtain a detectable signal when 2x the initial concentration of 2A12 or 4x the initial concentration of A05 was applied as the primary antibody (Supplementary Figure 3), Since this relates to the different affinities we determined for each clone using SPR, it appears that the native PAGE Western blots have less sensitivity than the SPR assays to detect the interactions between our VHH candidates and recombinant proteins.

## 4. Discussion

The therapeutic potential of VHHs has already been explored for other known tumour viruses in certain contexts. For example, VHHs against the S domain of the hepatitis B virus (HBV) envelope proteins (HBsAg) inhibited viral particle secretion in a mouse model (Serruys *et al*. 2009) and VHHs against the HBV nucleocapsid core protein (HBcAg) disrupted its subcellular localization (Serruys *et al*. 2010). However, the oncogenic influence of these proteins following HBV integration remains poorly understood (Tu *et al*. 2017) and the potential benefit of these VHHs in such a scenario is yet to be determined. In addition, VHHs against the hepatitis C virus (HCV) RNA-dependent RNA polymerase (NS5B) (Thueng-in *et al*. 2012), helicase (NS3 C-terminus) (Phalaphol *et al*. 2013), serine protease (NS3/4A) (Jittavisutthikul *et al*. 2015), membrane web and replication complex formation protein (NS4B) (Glab-Ampai *et al*. 2016), and envelope glycoprotein (E2) (Tarr *et al*. 2013) have been shown to inhibit viral replication or cell-to-cell transmission *in vitro*.

Thus far, for HPV16, only VHHs against the major capsid protein (L1) have been reported (Minaeian *et al*. 2012a and Minaeian *et al*. 2012b). Although these VHHs would serve the most benefit in diagnostic or prophylactic applications, there remained an unmet need for the development of therapeutic VHHs targeting the E6 oncoprotein, which we sought to address with this study. As target antigen properties such as solubility have been shown to influence the outcome of single-domain antibody library screens (Pardon *et al*. 2014, Henry and Tanha 2018) and recombinant HPV16 E6 is naturally prone to aggregation (Suarez and Travé 2018), it was initially anticipated that both immunizing with and panning against the solubility-enhanced His6MBP-E6 proteins would provide the best odds of isolating VHHs against native E6 protein epitopes. However, 20 of the 26 VHHs indicated by phage ELISA to be potential E6 binders had CDR3 sequences which were identified only in Library #1 (Supplementary Table 1), suggesting that immunizing with His6-GenScript E6 and panning the corresponding library against the solubility-enhanced His6MBP-E6 proteins was, instead, a more effective strategy. Although a subtractive panning technique was implemented, the majority of VHHs isolated from Library #2 were MBP binders. Perhaps the immune response to the His6MBP-E6 proteins was skewed more towards the larger MBP fusion partner (~42 kDa) than the E6 protein (~18 kDa) and further panning optimization will be required to facilitate the successful isolation of E6-binding VHHs from Library #2.

Of the 21 clones which were expressed as soluble VHHs and further characterized, we have thus far identified that A05, 2A12, and 2A17 bind recombinant HPV16 E6 with affinities in the nanomolar range. It is also possible that more rigorous screening of the remaining candidates using a better-suited ELISA format *(e.g.*, incubation of the target antigens with biotinylated VHHs followed by the application of HRP-conjugated streptavidin) or dot blots with increasing concentrations of VHH applied as the primary antibody will confirm additional clones as recombinant E6 binders. Further investigation will then be needed to determine whether these VHHs will similarly bind endogenous E6 protein *(e.g.*, in HPV16-positive biological samples). This is currently being tested for A05, 2A12, and 2A17, using an immunoprecipitation-based assay and cell lysates from HPV16-positive cell lines.

It will also be imperative to characterize what functional downstream effects they will elicit. One of the key protein-protein interactions which would be therapeutically beneficial to disrupt is that between E6 and the hijacked cellular E3A ubiquitin-protein ligase E6AP which, most notably, leads to viral-induced degradation of the tumour suppressor protein p53 (Martinez-Zapien *et al*. 2016). Loss of p53 interferes with the ability of infected cells to respond to a range of stressors including DNA damage, hyperproliferation, hypoxia, and oxidation (as reviewed by Bieging *et al*. 2014) and is thought to prevent cell cycle modulation by the HPV16 E7 protein from inducing apoptosis (Hoppe-Seyler *et al*. 2018). Such an approach has been favored by others who have attempted to functionally inhibit the HPV16 E6 protein using scFvs (Griffin *et al*. 2006, Lagrange *et al*. 2007, Amici *et al*. 2016). Molecules derived from sources other than natural immune system components have additionally been investigated and include zinc-ejecting compounds (Beerheide *et al*. 1999, Beerheide *et al*. 2000), small compounds (Baleja *et al*. 2006, Cherry *et al*. 2013, Malecka *et al*. 2014), inhibitory peptides (Liu *et al*. 2004, Sterlinko Grm *et al*. 2004, Dymalla *et al*. 2009, Stutz *et al*. 2015, Zanier *et al*. 2014), peptide aptamers (Butz *et al*. 2000), as well as bivalent inhibitors (Karlsson *et al*. 2015, Ramirez *et al*. 2015). VHHs targeting the C-terminus of the E6 protein are also of therapeutic interest for their potential to disrupt interactions with various host PDZ-domain containing proteins such as those involved in cell polarity and cell-cell signalling (as reviewed by Klingelhutz and Roman 2012). The utility of this concept was similarly demonstrated in a study by Belyaeva *et al*. 2014 using RNA aptamers. Hence, future work is needed to determine which E6 terminus is targeted by each of our VHH candidates and whether they can be used in combination to target both termini at once.

The previously reported E6-inhibitory molecules have had varying levels of success in decreasing cell proliferation and inducing apoptosis while maintaining minimal non-specific effects on HPV-negative cells and, notably, only one study reported *in vivo* data (Amici *et al*. 2016). In addition, therapeutic responses did not always occur in the manner expected. For example, although the solubility-improved 1F4 scFv (adapted from the 1F1 and 6F4 mAbs (Giovane *et al*. 1999)) was able to induce apoptosis when expressed as an intrabody in HPV16-positive cell cultures, it also non-specifically decreased proliferation in HPV-negative cells and the apoptotic pathway involved was not fully elucidated (Lagrange *et al*. 2007). The recent intrabody study characterizing the NLS-conjugated 17nuc scFv, which was isolated by Intracellular Antibody Capture Technology, also reported a necrotic response both *in vitro* and *in vivo*, rather than apoptosis (Amici *et al*. 2016). Interestingly, the E6/E6AP interaction-inhibiting compounds studied by Malecka *et al*. 2014 did not reduce the proliferation of the HPV16-positive cell line SiHa, despite restoring p53 protein levels, but did in HPV-negative PA-1 cells transfected with HPV16 E6. Such discord between p53 restoration and the subsequent induction of apoptosis in metastatic cervical carcinoma-derived HPV16 cell lines has also been discussed in the context of E6 suppression by siRNA (Togtema *et al*. 2018), leading us as well as others (Malecka *et al*. 2014) to question whether decades of *in vitro* culture have introduced subsequent mutations in apoptotic pathways which reduce reliance on continuous oncogene expression. Until it can be reliably demonstrated that CRISPR/Cas9 disruption of E6 at the DNA level (Stone *et al*. 2016) induces apoptosis in such cell lines without disruption to other host genes, it will be difficult to discern whether this is indeed the case and to what extent it may apply to freshly-derived patient samples. Nevertheless, p53 restoration in the absence of apoptosis may still provide therapeutic benefits, due to the additional roles this protein plays in regulating immune response (Muñoz-Fontela *et al*. 2016) and in inducing cellular senescence (Bieging *et al*. 2014). Although, senescent cells *(i.e.*, cells which exhibit permanent growth arrest but remain viable) secrete proinflammatory molecules such as cytokines and their accumulation in large numbers may undesirably counteract tumour regression (Coppé *et al*. 2008, Demaria *et al*. 2017).

A recent study by Stevanović *et al*. (2017) unexpectedly found that within populations of infused T cells which were cultured to enhance reactivity against the viral E6 and E7 proteins, it was tumour-infiltrating lymphocytes targeting neoantigens which played a key role in the regression of patients with metastatic HPV-positive cervical cancer. Although this study presents the results of only two patients (one HPV16-positive and one HPV18-positive) it indicates that added utility may be derived from simultaneously administering VHHs isolated against the viral E6 antigen together with those isolated against such patient-specific, non-viral targets. Their inclusion would also further increase the personalized aspect of our proposed therapeutic approach. With neoantigen expression vastly restricted to HPV-infected cells, such combinatorial treatment would still be anticipated to minimally affect surrounding healthy tissues. However, the timespan required to complete the VHH isolation and characterization process for each individual patient would be prohibitive, potentially restricting the applicability of this combinatorial treatment approach to more severe lesions.

With these concepts in mind, the further functional characterization of our E6 VHHs in monolayer cell cultures, three-dimensional raft cultures, as well as small animal models (as similarly described by Togtema *et al*. (2018) for siRNA) can be expected to identify their most optimal mode of application. The development of effective, clinically-compatible strategies for the intracellular delivery of the soluble VHHs themselves will also be required. Various antibody delivery techniques have been summarized by others (Steeland *et al*. 2016, Stewart *et al*. 2016, Böldicke 2017) and circumvent the patient risks associated with intrabody expression plasmids. In particular, several studies have demonstrated success in linking VHHs to the cell penetrating peptide, penetratin (Thueng-in *et al*. 2012, Phalaphol *et al*. 2013, Jittavisutthikul *et al*. 2015, Glab-Ampai *et al*. 2016). Our group has also explored the use of a high intensity focused ultrasound (HIFU)-based sonoporation technique to deliver HPV16 E6 mAbs (Togtema *et al*. 2012). It works by creating temporary pores in the cell membrane through HIFU-induced microbubble cavitation and can localize the delivery of therapeutic molecules to specific regions of target tissue (Phenix *et al*. 2014). Ideally, these approaches will need to be compared to determine which is most suitable in this context.

In addition to therapeutic limitations, researchers have also encountered difficulties using commercial mAbs for the immunodetection of HPV16 E6 and instead often examine downstream protein levels *(e.g.*, p53) as proxies (Bai *et al*. 2006, Gu *et al*. 2006, Yamato *et al*. 2008, Bousarghin *et al*. 2009, Tan *et al*. 2012). Immunocytochemistry has been particularly challenging (Jackson *et al*. 2013, Stutz *et al*. 2015), with limited success requiring the application of very specific staining protocols and complex image thresholding techniques (Jackson *et al*. 2013). However, as reviewed by Beghein and Gettemans (2017), the stability and antigen accessibility of VHHs have already facilitated their use for the immunofluorescent detection of various cellular antigens including vimentin (Maier *et al*. 2015), β-catenin (Braun *et al*. 2016), histone H2A-H2B heterodimer (Jullien *et al*. 2016), Vγ9Vδ2 T-cell receptor (de Bruin *et al*. 2016), and leukocyte receptor ChemR23 (Peyrassol *et al*. 2016). Accordingly, it can be anticipated that our VHH candidates may also demonstrate utility in these diagnostic types of applications. Thus, we ultimately envision the isolation of HPV16-E6 specific VHHs as a key first step, with the full range of their potential applications compared to that of traditional mAbs and fragments thereof to be fully realized in subsequent investigations.

## Author Contributions

The project was initially conceived by IZ and further developed with MT to then include JT and GH for their expertise in single-domain antibody research and production. MT, GH, GD, MRT and SR conducted the experiments, based on individual skills. Data compilation and analyses, as well as figure preparation was completed by MT with additional help from GH. MT wrote the manuscript, with substantial input from IZ, GH and JT and contributions from GD and MRT. All authors have participated in the critical revision of this manuscript and have approved the final version.

### Acknowledgements

The authors would like to acknowledge Shannon Ryan for her technical laboratory assistance and Robert Jackson for his critical input during the manuscript writing and revision process.

This work was supported by a Collaborative Health Research Projects (CHRP) grant between the Canadian Institutes of Health Research (CIHR) and the Natural Sciences and Engineering Research Council of Canada (NSERC) to IZ (CIHR: CPG 140188 and NSERC: CHRP 478520-15), an NSERC graduate student scholarship to MT (PGS-D3: 460717-2014), as well as the National Research Council Canada.

---

**Supplementary Figure 1.**
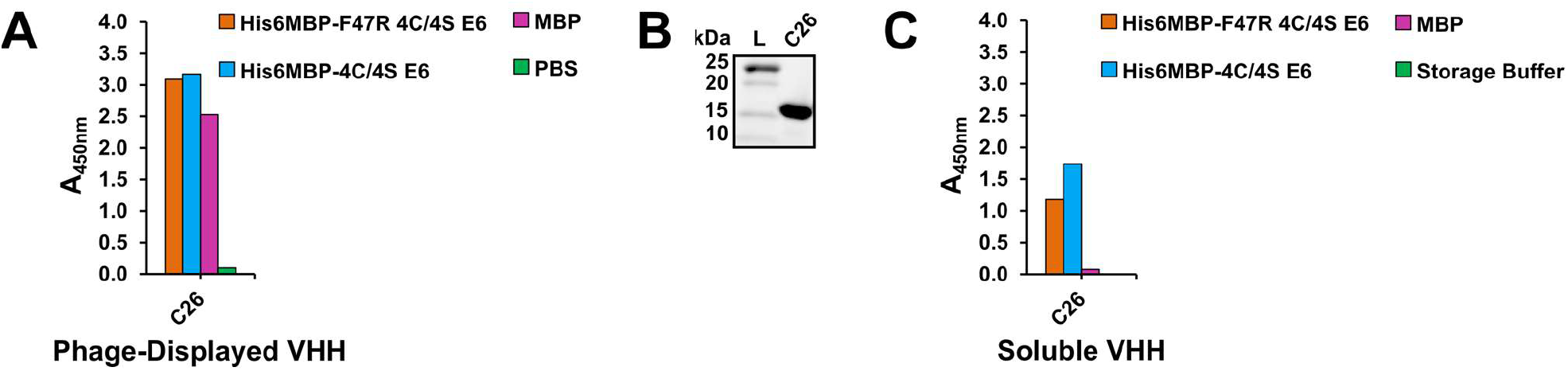
Clone C26 data. **(A)** Phage ELISA indicated that clone C26 was an MBP-binder, as phage displaying that VHH demonstrated affinity for both His6MBP-E6 proteins as well as MBP. Clones beginning with “C” were isolated from Library #2 round 1 eluted phage titer plates. (B) Reducing SDS-PAGE was used to analyze the purity of the VHH following IMAC elution. Approximately 3 μg of sample was loaded on the gel. (C) An ELISA using 100 μg/mL of clone C26 demonstrated that this purified VHH was functional and maintained its MBP-binding characteristics.

**Supplementary Figure 2.**
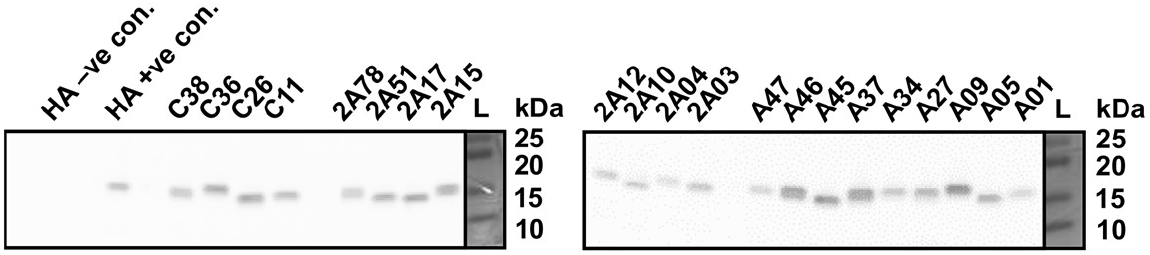
Confirmation that the HA tags remained intact on the purified VHHs. *Reducing SDS-PAGE Western blots using an anti-HA + HRP antibody demonstrated the HA tags had not been degraded on the purified VHHs which were stored long-term at −20 °C. Approximately 5 ng/lane VHH was loaded. The HA positive (+ve) control is a VHH with both HA and His6 tags which was isolated against an unrelated target antigen. The HA negative (-ve) control is a VHH which alternatively has both Myc and His6 tags*.

**Supplementary Figure 3.**
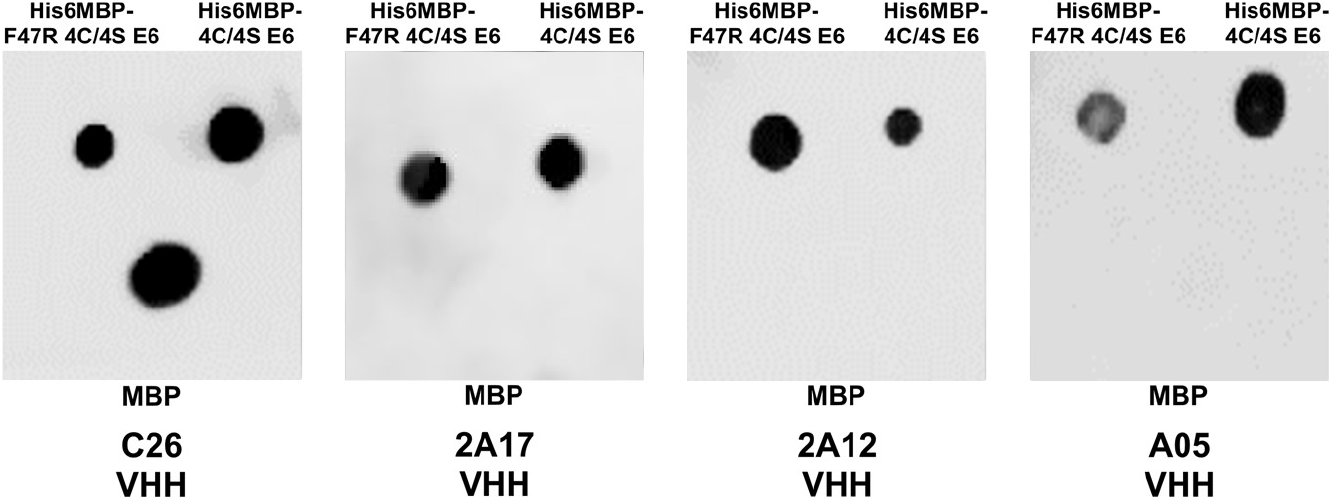
Dot blots testing increased VHH concentrations. As observed with the native PAGE Western blots, incubation of the dot blot membranes with 2.7 μg/mL VHHs C26 or 2A17 as the primary antibody yielded a detectable signal. However, 2x this concentration (5.4 μg/mL) and 4x this concentration (10.8 μg/mL) had to be applied to obtain a detectable signal with VHHs 2A12 and A05, respectively. Approximately 2 μg of His6MBP-F47R 4C/4S E6, His6MBP-4C/4S E6, and MBP was spotted on the membrane for each dot.

**Supplementary Table 1.** Eluted VHHs with the same complementarity-determining region 3 (CDR3) sequences.

| Clone | Additional clones with same CDR3 sequence |  |  |  | Total occurrences of each CDR3 sequence |
| --- | --- | --- | --- | --- | --- |
|  | Library #1, round 1 | Library #1, round 2 | Library #2, round 1 | Library #2, round 2 |  |
| A01 | - | - | - | - | 1 |
| A05 | - | - | - | - | 1 |
| A09 | A13 | 2A34 | - | - | 3 |
| A24 | - | - | - | - | 1 |
| A26 | - | - | - | - | 1 |
| A27 | - | - | - | - | 1 |
| A34 | - | 2A67, 2A69 | - | - | 3 |
| A37 | - | - | - | - | 1 |
| A45 | - | - | - | - | 1 |
| A46 | - | - | - | - | 1 |
| A47 | - | 2A05, 2A46 | C41 | - | 4 |
| C06 | - | - | - | - | 1 |
| C11 | - | 2A08, 2A13, 2A72, 2A79, 2A89 | C22 | - | 7 |
| C27 | - | - | - | - | 1 |
| C36 | - | - | - | 2C37 | 2 |
| C38 | - | 2A18, 2A26, 2A33, 2A39 | - | - | 5 |
| 2A03 | - | 2A37, 2A62 | - | - | 3 |
| 2A04 | - | - | - | - | 1 |
| 2A10 | - | - | - | - | 1 |
| 2A12 | - | 2A45 | - | - | 2 |
| 2A15 | - | - | - | - | 1 |
| 2A17 | - | - | - | - | 1 |
| 2A51 | - | - | - | - | 1 |
| 2A55 | - | - | - | - | 1 |
| 2A78 | - | 2A06, 2A48, 2A63, 2A66 | - | - | 5 |
| 2A90 | - | - | - | - | 1 |

